# A Purinergic Mechanism Underlying Metformin Regulation of Hyperglycemia

**DOI:** 10.1101/2022.09.27.509754

**Authors:** J. Senfeld, Q. Peng, Y. Shi, S. Qian, J. Shen

## Abstract

Metformin, created in 1922, has been the first-line medication treating type 2 diabetes mellitus for almost 70 years; however, its mechanism of action has been heavily debated, partly because most prior studies used supratherapeutic concentrations exceeding 1mM. Here we report that at a clinically relevant concentration of 10 μM, metformin blocks high glucose-stimulated ATP secretion from hepatocytes mediating its antihyperglycemic action. Following glucose administration, mice demonstrate increased circulating ATP concentrations, which are prevented by metformin. Extracellular ATP through P2Y2 receptors (P2Y2R) compromises insulin-induced AKT activation and increases hepatic glucose production. In addition, metformin-dependent improvement in glucose tolerance is abolished in P2Y2R-null mice. Thus, removing the target of extracellular ATP, P2Y2R, mimics the effects of metformin, revealing a novel purinergic antidiabetic mechanism for metformin.

**One-Sentence Summary:** Metformin abolishes high glucose-stimulated ATP secretion, preventing purinergic mechanism-mediated hepatic insulin resistance and glucose production.

## Main Text

The global prevalence of obesity and Type 2 Diabetes Mellitus (T2DM) surmounts to epidemic proportions and is expected to afflict 700 million people by 2045 (*1*). Generated in 1922 (*2*) and used for almost 70 years, metformin is a first-line pharmacotherapy taken by over 150 million patients worldwide to improve glycemic control. Metformin effectively suppresses hepatic glucose production and enhances insulin sensitivity (*3*); however, its mechanism of action (MOA) remains enigmatic. It has been reported that metformin inhibits mitochondrial complex I, disrupting respiration at doses exceeding therapeutically achievable concentrations (*4, 5*). Another documented MOA was that metformin stimulates AMPK, which has recently been brought into question because metformin’s antidiabetic capabilities were unaffected in liver-specific AMPK knockout mice (*6*). We also found that metformin up to 30μM neither changed intracellular ATP levels nor activated AMPK in human hepatocytes (fig. S1). Further, forced expression of gluconeogenic genes, including glucose-6-phosphatase, failed to affect the suppression of hepatic gluconeogenesis by metformin (*6*). Therefore, we may reasonably conclude that metformin operates, in part, in an AMPK- and transcriptional-independent manner. Unlike other antidiabetic medicines, metformin seldom affects normal fasting blood glucose levels leading us to consider that metformin’s MOA may rely on physiological events initiated during increased blood glucose concentrations. Thus, we examined the hyperglycemia-induced MOA of metformin.

Extracellular high glucose is a chemical stressor, inducing ATP release from various metabolic tissues, including the liver (*7, 8*). Thus, we proposed that glucose-stimulated ATP secretion (GSAS) represents an autocrine and paracrine mechanism that insulin-sensitive tissues employ to control glucose homeostasis through the purinergic signaling axis. Figure 1A shows that challenging HepG2 human hepatocytes with 25 mM glucose, but not by 25mM mannitol, induced time-dependent ATP secretion, indicative of a specific GSAS in hepatocytes. Interestingly, co-treatment of the cells with 30 μM metformin abolished GSAS in all tested time points (Fig. 1A), which was not due to a change in extracellular glucose level (fig. S2). Further studies indicated that the effective concentration of metformin to block GSAS is around 10 μM (Fig. 1B), consistent with the known clinical blood concentration range of 10 ~ 40 μM for metformin (*9*). Importantly, this GSAS-blocking effect was fully mimicked by phenformin and buformin at a much higher potency and fully replicated in primary human hepatocytes (fig. S3), consistent with their clinical dose profiles. In contrast, proguanil, an analog of metformin with no antidiabetic effect, had no inhibition on GSAS (Fig. 1C-E). To complement our luciferase ATP assays not interfered with by metformin at 30 μM (fig. S4), we generated a HepG2 cell line in which the cell membrane stably expressed a fluorescent sensor detecting extracellular ATP (iATPSnFR). Figure 1J shows that compared with control 5.5 mM glucose, treatment of HepG2-iATPSnFR cells with 25 mM glucose for 60 minutes exhibited more green fluorescent signals co-localized with the cell membrane red fluorescence. Co-treatment of the cells with 30 μM metformin abolished the green fluorescence, indicating that metformin suppresses GSAS (Fig. 1J and fig. S5), which was confirmed by a different fluorometric ATP quantification assay (fig. S6).

**Fig. 1.**
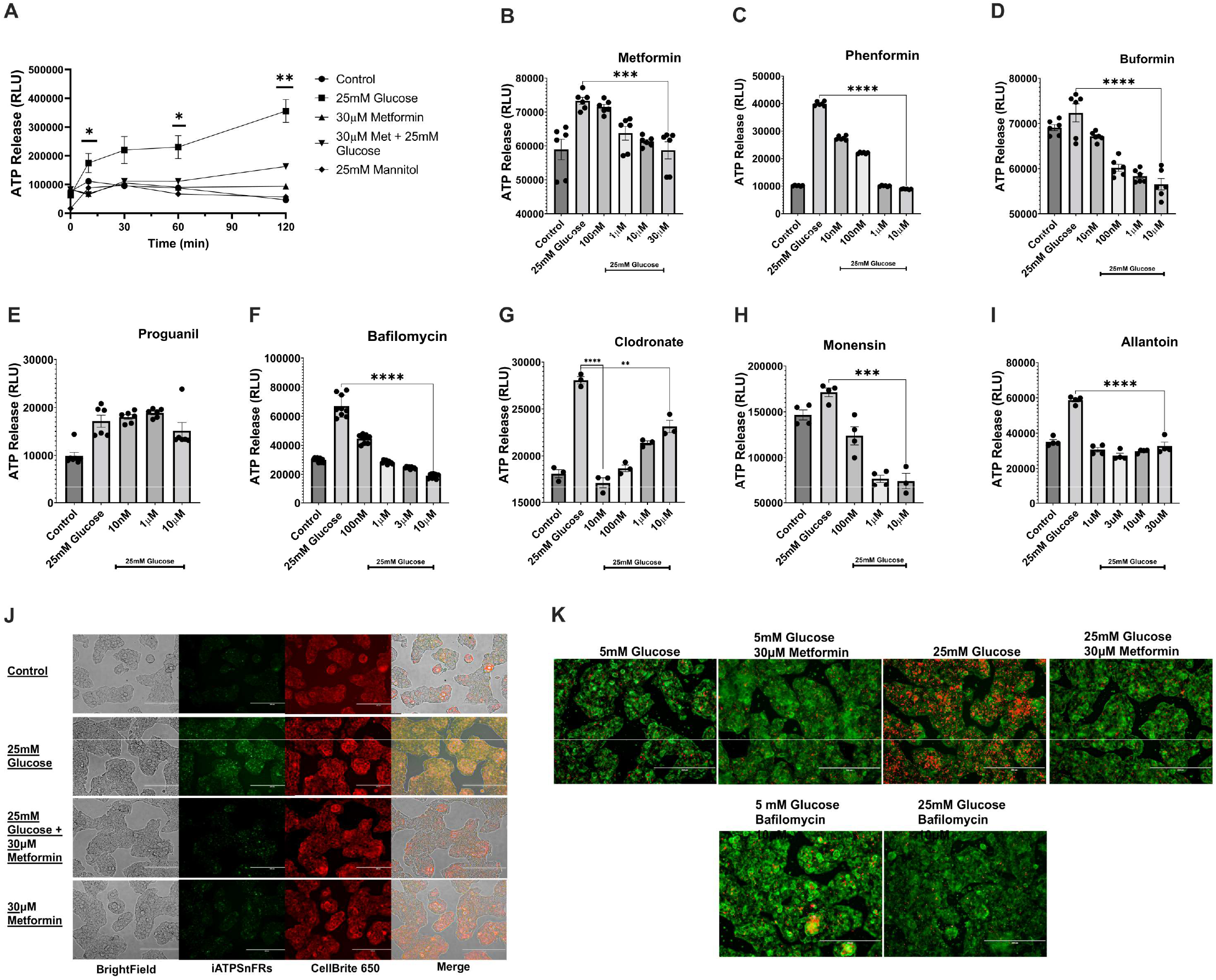
Metformin, alongside other biguanides and endolysosomal alkalizing agents, prevents glucose-stimulated ATP secretion (GSAS). (**A**) Time course observation of ATP secretion originating from HepG2 hepatocytes treated with 5mM, 25mM glucose, or 25mM mannitol with or without co-treatment with 30μM metformin. *P<0.05, **P<0.01. (**B-I**) Dose-dependent effects on GSAS by antidiabetic biguanides metformin, phenformin, buformin, antimalaria biguanide proguanil, endolysosomal targeting agents bafilomycin, clodronate, monensin, and ATP metabolite allantoin. Data are means ± SEM of three independent experiments. ***P<0.005, ****P<0.001. (**J**) Detection of extracellular ATP signal (green) by HepG2 cells transfected with iATPSnFR, a plasma membrane-bound ATP sensor, was observed following the indicated treatments and co-localized with a cell membrane red dye Cellbrite650 (yellow). (**K**) Effect of metformin and high glucose treatment on endolysosomal pH change monitored by LysoView633 (red), a pH-sensitive endolysosomal probe. CellBrite488 (green) was used for plasma membrane staining and bafilomycin as a positive control. Shown are representative of three independent experiments. Scale bar: 200μm

We then explored the origin of ATP released in response to hyperglycemia. Curiously, metformin’s primary site of action, the liver, expresses significantly more vesicular nucleotide transporters (VNUT) mRNA than any other tissue in the human body (*10*), and we confirmed its cytoplasmic expression in HepG2 cells (fig. S7). VNUT-knockout mice demonstrated improved glucose tolerance and insulin sensitivity (*11, 12*). Further, VNUT activity is coupled to the action of V-type-ATPase that has recently been implicated in the MOA of metformin (*9*). Therefore, we first considered a vesicular origin of GSAS. To evaluate the contribution of the endolysosomal system, we pharmacologically depleted endolysosome ATP content by disruption of the proton gradient or inhibition of ATP transport. Pretreatment of the cells with clodronate or bafilomycin, VNUT and V-type-ATPase inhibitors, respectively, produced a dose-dependent reduction in GSAS (Fig. 1F-G). Consistent with this, pretreatment of the cells with monesin, a chemical mimetic of Na^+^/H^+^ exchanger known to alkalize luminal pH of endolysosomes, reduced GSAS (Fig. 1H). In addition, we verified that high glucose treatment acidified endolysosomes, as evidenced by increased LysoView pH-dependent probe staining, which was suppressed by 30μM metformin (Fig. 1K). Thus, our results support the notion that metformin may interfere with the proton gradient of the endolysosome, leading to the depletion of vesicular ATP content, which was further supported by our observation that metformin prevented MANT-ATP accumulation (fig. S8). We then hypothesized that a metabolite of the purine degradation pathway might serve as an endogenous feedback mechanism to control GSAS. Allantoin, a metabolite of uric acid negatively correlated with T2DM (*13, 14*), displayed potent inhibition of GSAS at physiological concentrations (Fig. 1I). However, humans harbor a loss-of-function mutation in the gene encoding uric oxidase; consequently, humans cannot convert uric acid to allantoin enzymatically, potentially predisposing modern humans to diabetes.

Next, we aimed to define how extracellular ATP affects hepatocyte control of glucose homeostasis, focusing on the nucleotide P2Y receptors since the adipocyte P2Y6 and P2Y14 receptors are implicated in metabolic disease (*15, 16*). RT-PCR analysis indicated that P2Y2 and P2Y11 are the predominant ATP-sensitive P2Y receptors expressed in HepG2 cells (Fig. 2A). Ca^2+^ mobilization assays showed that both ATP and UTP induced a dose-dependent increase of intracellular Ca^2+^, which were abolished by ARC-118925XX, a P2Y2R-selective antagonist (Fig. 2B-D), and no functional P2Y6R and P2Y11R were detected (fig.S9). siRNA silencing of the P2Y2R also abolished ATP- or UTP-induced Ca^2+^ signaling (Fig. 2E), supporting P2Y2R as the primary ATP- and UTP-sensitive P2Y receptor in hepatocytes. We then explored P2Y2R-mediated other signaling pathways. Figure 2F & fig. S10 show that stimulation of the P2Y2R activated the JNK and p38 pathways with slight activation of the ERK1/2 and no activation of the AKT pathways. Activation of JNK and p38 pathways indicates an inflammatory response, a condition often preceding metabolic dyshomeostasis and insulin resistance (*20–22*). Further, the apparent decrease of basal AKT phosphorylation by ATP and UTP made us hypothesize that the P2Y2R may have an inhibitory effect on the AKT pathway. This was supported by our finding that ATP and UTP dose-dependently suppressed insulin-induced AKT phosphorylations at the T308 and S473 sites (Fig. 2G-H) without compromising the ERK1/2 pathway (fig.S11). The expression and function of P2Y2R in control of insulin-AKT signaling were also confirmed in primary human hepatocytes (fig. S12 & S13). To further explore the underlying mechanism(s) responsible for P2Y2R-mediated AKT inhibition, we hypothesized that depletion of PIP_2_, a substrate shared by PI3K and PLC, may be involved. Figure 2I shows that activation of the P2Y2R decreased insulin-induced AKT phosphorylation, which was entirely prevented by U73122, a specific PLC inhibitor. In addition, stimulation of the P2Y2R suppressed insulin-induced PIP_3_ generation (fig. S14), and supplementation of PIP_2_ at least partially rescued P2Y2R-mediated AKT inhibition (fig. S15). These results indicate that P2Y2R selectively impairs insulin-induced AKT signaling, consistent with numerous reports that insulin resistance and T2DM are linked to a disruption of a branch of insulin receptor signaling. Further, identifying P2Y2R as the anti-insulin signaling mediator advances early discoveries demonstrating potent inhibition of insulin signaling and hepatocyte glucose production by extracellular nucleotides (*17, 23–30*). All these shreds of evidence support our hypothesis that metformin improves hepatic insulin sensitivity by suppressing ATP release and signaling, consistent with a previous report utilizing insulin receptor-null mice concluding “for metformin to have any effect, an operating insulin-signaling system in the liver is mandatory” (*31*).

**Fig. 2.**
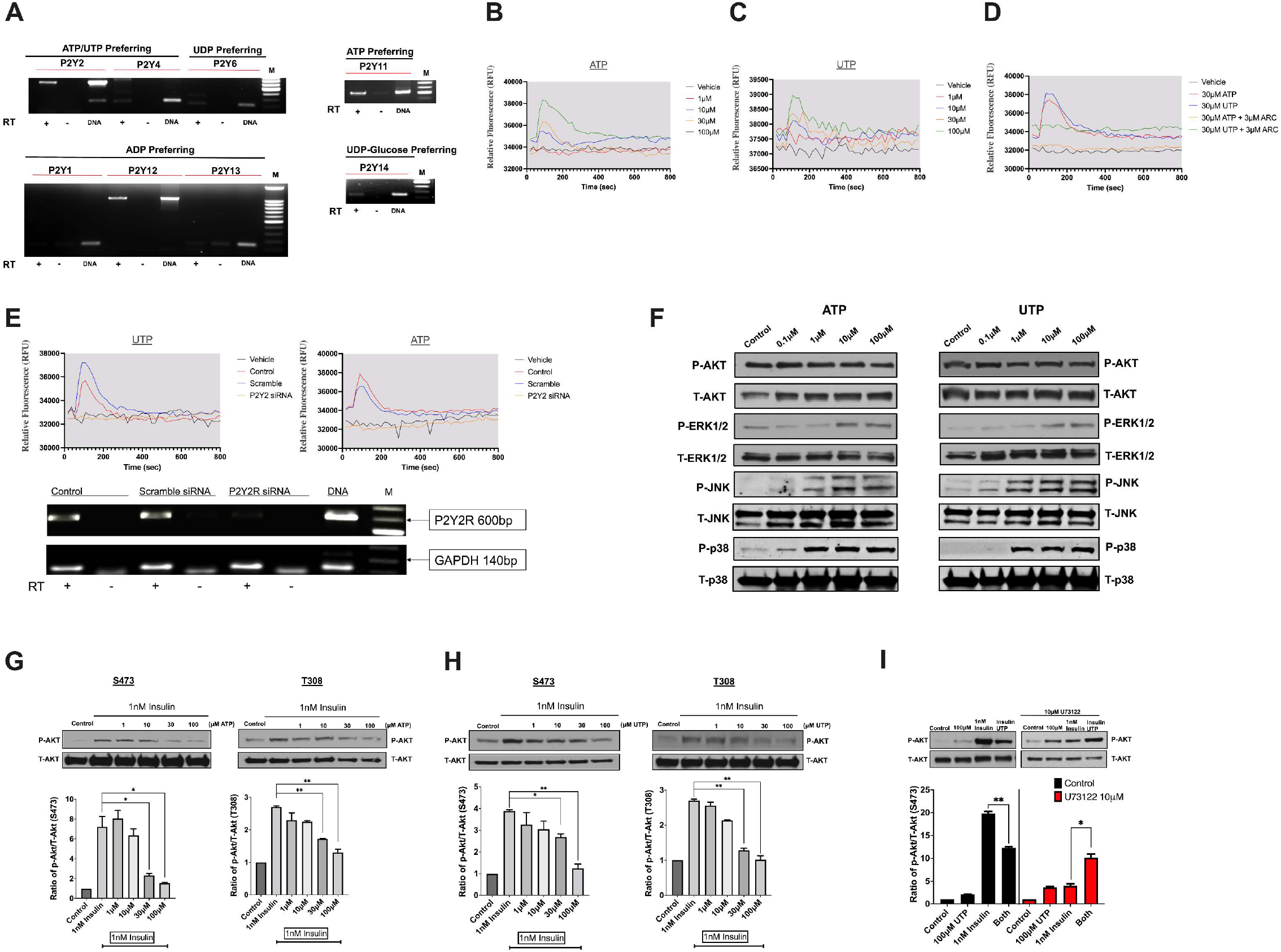
Identification of AKT-inhibitory purinergic P2Y2R in hepatocytes. (**A**) RT-PCR analysis of mRNA expression profile of the eight known P2Y receptor subtypes in HepG2 cells. (**B, C**) HepG2 cells were stimulated with P2Y2R agonists ATP and UTP at the indicated concentrations, and [Ca^2+^]_i_ mobilization was monitored fluorometrically. (**D**) Effect of P2Y2R-selective antagonist ARC-118925XX (3μM, 45 min) on nucleotide-induced [Ca^2+^]_i_ mobilization. (**E**) Hepatocytes were pretreated for 48 hr with P2Y2R siRNA (50nM) or scrambled siRNA (50nM), and nucleotide-induced [Ca^2+^]_i_ mobilization was assessed. P2Y2R knockdown was confirmed through RT-PCR. (**F**) Western blotting was employed to determine P2Y2R signaling to the MAPK and AKT pathways following a 10 min stimulation by ATP or UTP at the indicated doses. (**G, H**) Effect of P2Y2R stimulation on insulin-induced AKT phosphorylations at the sites of T308 and S473. (**I**) Effect of PLC-selective inhibitor U73122 (10μM, 45min) on P2Y2R-mediated suppression of insulin-AKT signaling. The data shown are representative of three independent experiments. *P<0.05, **P<0.01

We then assessed whether P2Y2R affects hepatocyte glucose production (HGP). Unlike skeletal muscle and adipose tissues, the hepatic glucose transporter, GLUT2 activity is not dependent on insulin signaling, enabling the liver to take in and release glucose bidirectionally (*17*) during the fed and fasted state. It was reported that intravenous administration of adenine nucleotides stimulates HGP (*18–21*). However, a nucleotide receptor controlling HGP remained elusive; therefore, we first stimulated hepatocyte P2Y2R with ATP and UTP, finding a dosedependent reduction in glucose uptake (Fig. 3A-B), suggesting a possible glucose release in response to P2Y2R activation. We then monitored gluconeogenesis to determine if decreased glucose uptake was correlated to increased glucose production. Figure 3C shows that stimulating the HepG2 cells by ATP or UTP significantly increased gluconeogenesis. Notably, insulin failed to suppress nucleotides-induced gluconeogenesis (Fig. 3C). However, pretreatment with selective P2Y2R antagonist, ARC-118925XX, effectively prevented nucleotide-induced gluconeogenesis (Fig. 3C). Early studies found that perfusing murine livers with ATP or UTP increased glycogen phosphorylase activity and glycogenolysis (*22–24*). In line with these findings, we found that stimulation of the P2Y2R for 10 min or 45 min significantly decreased glycogen content in HepG2 cells, an effect comparable to that induced by a maximal dose of glucagon (Fig. 3D). In addition, P2Y2R activation inhibited insulin-induced phosphorylation of GSK3β (Fig. 3E-F), providing another layer of mechanistic understanding of P2Y2R control of insulin signaling and glycogen synthesis in hepatocytes as summarized in Fig. 3G.

**Fig. 3.**
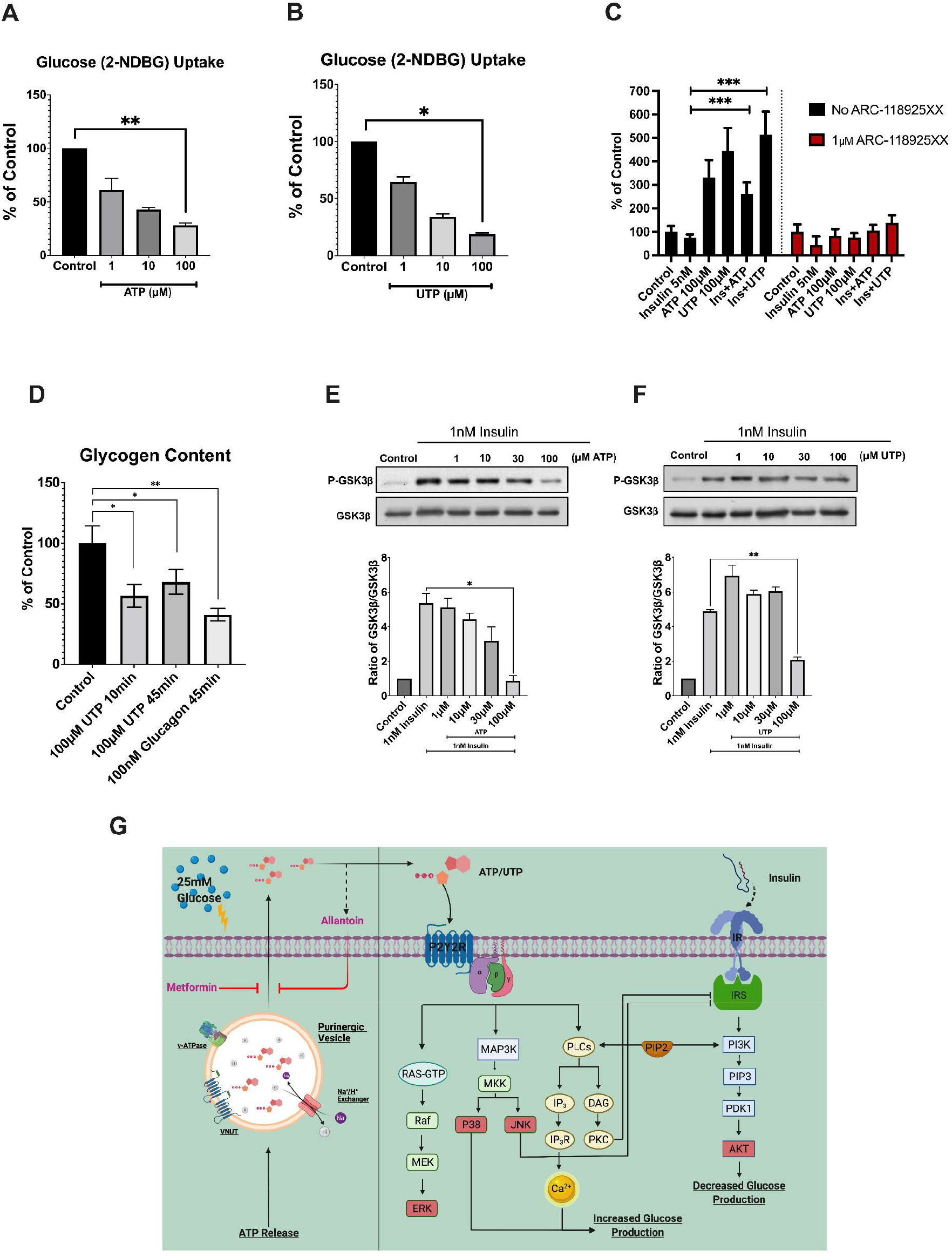
P2Y2R control of hepatocyte glucose production. (**A, B**) HepG2 cells were stimulated with ATP or UTP at the indicated concentrations for 20 min, then 2-NDBG (150 μg/mL) was added for 10 min before quantifying intracellular 2-NDBG. *P<0.05, **P<0.01, n=5. (**C**) Hepatocyte gluconeogenesis was measured by an Amplex red coupled reaction following a 3.5-hour simulation with ATP or UTP ((100μM) or insulin (1nM) using 10 mM sodium lactate as gluconeogenic substrate. All cells were pretreated for 45 min with ARL-67156 (100μM) to prevent nucleotide degradation and with or without ARC-118925XX (1μM) as indicated. ***P<0.005, n=5. (**D**) Hepatocyte glycogen content was measured fluorometrically following stimulation with UTP (100μM) or glucagon (100nM) for the indicated duration of time. *P<0.05, **P<0.01, n=5. (**E, F**) Insulin-induced inhibition of GSK3β by phosphorylation (S9) was assessed by Western blotting after 10 min co-stimulation with ATP or UTP and insulin at the indicated concentrations. *P<0.05, **P<0.01, n=5. (**G**) A diagram shows a new fine-tuning GSAS mechanism underlying P2Y2R-controlled insulin signaling, glucose production, and its pharmacological intervention by metformin in hepatocytes.

Finally, we determined whether metformin’s antihyperglycemic action *in vivo* is related to P2Y2R. Animal studies indicate that CD39 deletion results in a hepatic insulin-resistant phenotype in mice (*25*), and hyperglycemia decreases gene expression and activity of ectonucleotidases (*26*), suggesting that hyperglycemia may potentiate liver purinergic signaling. Indeed, circulating adenine nucleotides were elevated in T2DM patients (*27*), and intraperitoneal injection of AMP increased mouse blood ATP and glucose levels (*28*). In db/db mice, elevated plasma adenine nucleotides strongly correlated to increased plasma glucose levels (*29*). Prompted by these observations and our own in *vitro* findings, we determined whether glucose loading affects the blood level of ATP. Figure 4A shows that intraperitoneal administration of glucose significantly increased blood ATP levels at 30 min and 60 min, which were diminished considerably by metformin pretreatment at 50mg/kg (i.v.), a mode of administration and dose shown to reach mouse plasma metformin concentrations around 10~20 μM within the first hour (*30*). We then performed GTT and ITT assays in mice fed with a standard chow diet. Figure 4B shows that P2Y2R-null (KO) mice exhibited a better glucose tolerance compared with littermate-controlled wild-type (WT) mice, and this effect was mimicked by metformin treatment in WT but not in KO mice (Fig. 4C-D). Since a recent report indicated that mice lacking P2Y2R are resistant to dietinduced obesity and display improved glucose and insulin tolerance (*31*), we compared ITT profiles in mice fed with a high-fat diet versus a regular diet. Figure 4F shows that metformin treatment significantly improved insulin tolerance in WT mice fed a high-fat diet. In contrast, the same metformin treatment did not impact insulin sensitivity in WT mice fed a standard chow diet (Fig. 4E) and KO mice fed a high-fat diet (Fig. 4G). Further, we confirmed that liver P2Y2R and VNUT were upregulated in mice fed a high-fat diet compared to normal healthy mice (Fig. 4H). These data support our notion that metformin’s action relies on the presence of hyperglycemia and P2Y2R.

**Fig. 4.**
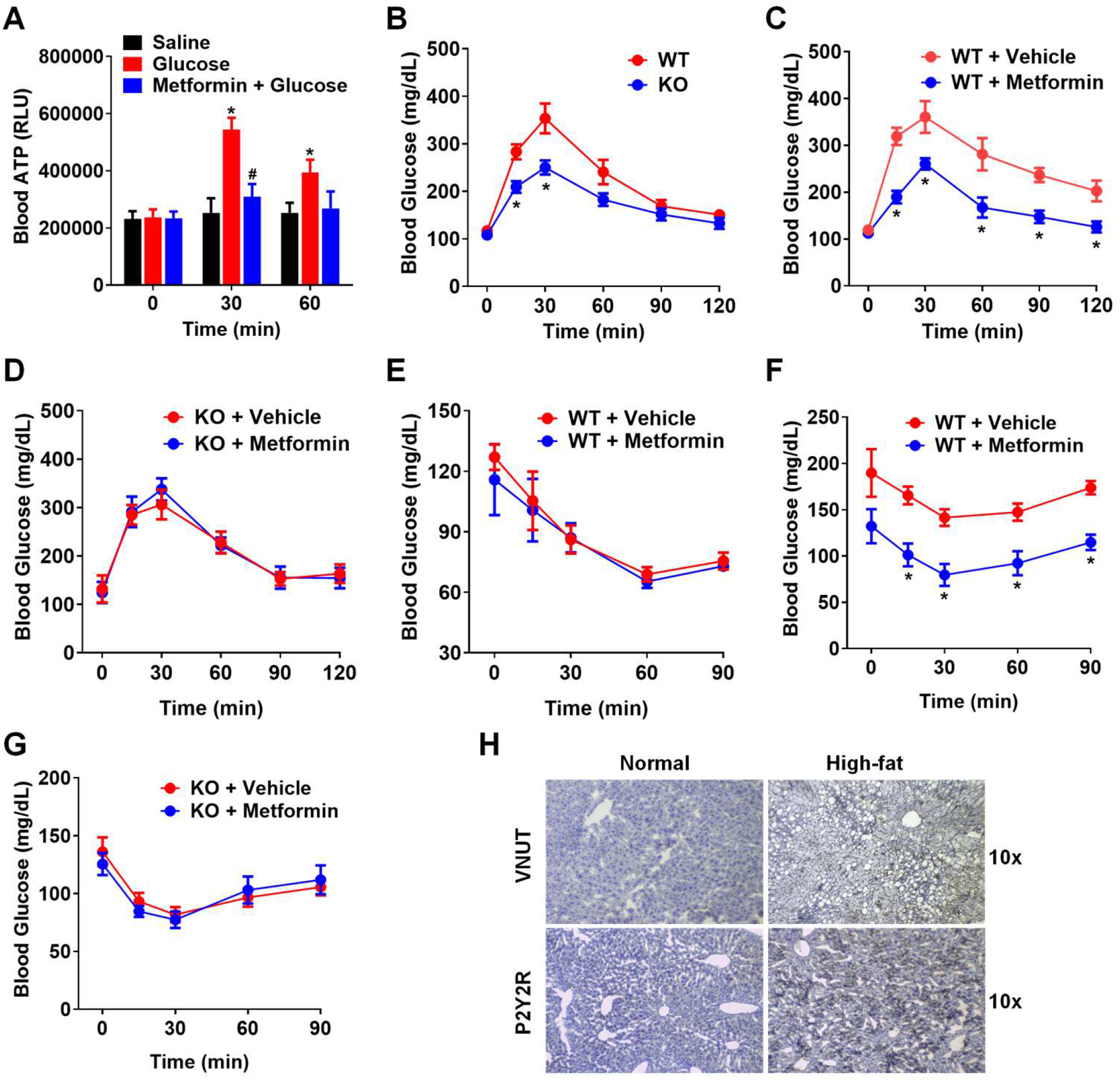
Hyperglycemia and P2Y2R dictate metformin’s anti-hyperglycemic action. (**A**) Effect of high glucose loading (1.5g glucose/kg, i.p.) on the relative levels of mouse blood ATP at the indicated times. WT mice were pretreated for 15 min with saline or metformin (50mg/kg, i.v.) through tail vein injection before glucose loading. n=6 mice in each group. *P<0.05 compared with the saline group; #P<0.05 compared with the glucose group. (**B-D**) Glucose tolerance test (GTT, 1.5g glucose/kg, i.p.) comparing P2Y2R-KO versus WT mice, metformin treatment (50mg/kg, i.v.) versus saline in WT or P2Y2R-KO mice on a standard chow diet. n=6~10, *P<0.05. (**E**) Insulin tolerance test (ITT, 0.75 IU insulin/kg, i.p.) comparing vehicle versus metformin-treated mice on a standard chow diet. n=6~9. (**F, G**) ITT was comparing saline versus metformin treatment on WT or P2Y2R-KO mice fed with a high-fat diet for 16 weeks. n=6~10, *P<0.05. (**H**) Immunohistochemical staining of VNUT and P2Y2R in liver sections of representative mice fed with normal chow or high-fat diet for 16 weeks.

We have demonstrated that metformin, alongside other antidiabetic biguanides, prevents hepatic GSAS and displays no antihyperglycemic effect in P2Y2R-null mice, indicating that metformin’s MOA involves disruption of the purinergic signaling axis. Hormones, glucagon and insulin have been exhaustively studied due to their paramount roles in systemic metabolic homeostasis; however, nucleotide release could represent a transient and local mechanism employed by insulin-sensitive tissues in response to nutrient overload operating synergistically and competitively towards glucagon and insulin, respectively. Normalizing pathological nucleotide release could clarify the beneficial phenotypes of metformin treatment spanning diabetes, obesity-induced inflammation, aging, and other metabolically rooted pathologies.

## Acknowledgments

We thank Dr. Peter Panizzi’s lab for assistance with cryostat liver tissue sectioning.

## Author contribution

J.Shen conceived and supervised the project. J.Senfeld performed most of the cell culture experiments, and QP performed all the in vivo mouse experiments. YS and SQ performed PIP_3_ ELISA and immunohistochemistry studies. J.Senfeld wrote the original draft, and QP and J.Shen edited the manuscript.

## Data and materials availability

All data are available in the main text or the supplementary materials. All plasmids, cell lines, and mice are available through a material transfer agreement with Auburn University by contacting the corresponding author.

## Supplementary Materials for

### Materials and Methods

#### Materials

Minimum Essential Medium (MEM) or Eagle’s Minimal Essential Medium (EMEM) were purchased from Lonza. FluoForteTM Kit was purchased from Enzo Life Sciences. Fetal Bovine Serum (FBS) was purchased from Thermo Fisher Scientific. DNA primers were purchased from Integrated DNA Technologies. DNase, RNeasy, and DNeasy kits were purchased from Qiagen. The cDNA synthesis kit was purchased from Applied Biosystems. Purified UTP and ATP were obtained from MilliporeSigma. HepG2 hepatocytes were purchased from ATCC, and Primary Human Hepatocytes (PHH) were purchased from Applied Biological Materials Inc.. Primary and secondary antibodies were obtained from Cell Signaling Technology and ABclonal. siRNA, ON-TARGET plus SMART pool L-003688-00-0005, human P2RY2, NM_002564, and DharmaFECT-1 transfection reagents were purchased from Dharmacon. The plasmid of the ATP sensor iATPSnFRs was purchased from Addgene. Metformin and Monensin were purchased from Enzo Life Science, phenformin was from BioVision, buformin was purchased from Cayman Chemical, Proguanil was from MilliporeSigma, and bafilomycin was purchased from BioViotica. All other reagents were obtained from Tocris.

#### Cell cultures

HepG2 cells were filtered by 70 μm cell strainer and grown in a monolayer, cultured in EMEM (also called MEM) supplemented with 10% FBS at 37 °C in a humidified atmosphere of 5% CO_2_. Primary human hepatocytes (PHH) were also cultured in EMEM supplemented with 10% FBS at 37 °C in a humidified atmosphere of 5% CO_2_. Before stimulation, cells were seeded to grow for 24 h, reaching ~80–90% confluence, and starved for 3 h, 6 h, or overnight. When inhibitors or antagonists were used, cells were pretreated with the inhibitor/antagonist for 35min to 1 h before cell stimulation by different agonists at the specified times and concentrations.

#### Intracellular Ca^2+^ mobilization assay

Cells were seeded at a density of 4×10^4^ per well into 96-well culture plates and cultured for one day. On day two, the original medium was removed. The assay medium from the FluoForte™ kit containing the Ca^2+^ dye was added, and receptor-mediated Ca^2+^ mobilization was determined as we previously described (*32*). Fluorescence was determined immediately after adding different reagents, with an excitation wavelength set to 490 nm and an emission wavelength set to 525 nm, and readings were taken every 1s for 640s. For the antagonist inhibition experiment, cells were pre-incubated with the antagonist for 45 min before agonist addition. Measurement of Ca^2+^ signal was performed with the fluorometer plate reader (B.M.G. FLUOstar), the results of which were shown as relative fluorescence units (RFU).

#### ATP secretion assays

##### Luminescent method

ATP secretion from hepatocytes was quantified utilizing ENLIGHTEN ATP Assay System Bioluminescence Detection Kit (Promega). Hepatocytes were cultured in a 6-well plate 24 h before experimentation and starved for 3-5 h before the start of the experiment. NTPDase inhibitor ARL67156 (Tocris) was added to every well at a concentration of 100μM for 30 min to inhibit the enzymes that degrade extracellular ATP. Compounds metformin, phenformin, buformin, proguanil, monensin, bafilomycin, clodronate, and allantoin were pretreated for 35 min at concentrations from 10 nM to 30 μM before supplementing 5- or 25 mM D-glucose. Measurement of extracellular ATP was accomplished by combining 100 μL assay reagent with 100 μL medium extracted from each well in a white 96-well plate. Quantification was completed in a Glomax 96 microplate luminometer (Promega), resulting in data being presented as relative light units (RLU).

##### Fluorometric method

ATP secretion from hepatocytes was also quantified utilizing a colorimetric/fluorometric ATP Assay Kit (Abcam). Hepatocytes were cultured in a 6-well plate for 24 h and then starved for 3-6 h before the start of the experiment. Additionally, NTPDase inhibitor ARL67156 was first added to every well at a concentration of 100 μM for 30 min. Compound metformin was pretreated for 35 min at concentrations from 1 μM to 30 μM before supplementing 5- or 25 mM D-glucose. Extracellular ATP was measured by combining 50 μL assay reagent with 50 μL medium extracted from each well in a black/transparent bottom 96- well plate (Greiner), incubated in darkness at room temperature for 30 min. Data was collected from a 96-well plate reader using Ex/Em = 535/587nm and reported in relative fluorescent units (RFU).

#### Intracellular ATP quantification

Hepatocytes were plated in 6-well plates containing the growth medium at 80% confluency. ATP quantity in hepatocytes was determined utilizing a colorimetric/fluorometric ATP Assay Kit (Abcam, ab83355). Before performing the ATP quantification assay, a fresh serum-free culture medium was added to every well for 3-6 h and then treated with 30 μM metformin or PBS for 2 h. Cell lysates were then collected using the Deproteinizing Sample Preparation Kit (Abcam, ab204708). ATP quantity in the cell lysates was determined based on the phosphorylation of glycerol to generate a product detectable by fluorometry or absorbance. Briefly, 50 μL assay reagent was mixed with 50 μL lysate in a transparent bottom 96-well plate and incubated in darkness at room temperature for 30 min. ATP concentration was then determined by measuring absorbance at OD 570 nm (colorimetric). The results are shown as absorbance (abs).

#### Measurement of the blood level of ATP

Mice were pretreated for 15 min with saline (0.9% NaCl) or metformin (50mg/kg, i.v.) through tail vein injection, after which 5 uLof tail blood was mixed with 45 uL saline to minimize platelet activation. The baseline level of ATP in the diluted blood sample was immediately measured by the luciferin-luciferase assay as previously described. Then, mice were injected intraperitoneally with D-glucose (1.5g/kg body weight) in normal saline (0.9% NaCl) or just saline as a control. Blood samples were further collected after 30 min and 60 min of D-glucose injection, and measurement of ATP was executed by combining 50 μL assay reagent with 50 μL diluted blood in a white 96-well plate. Quantification was accomplished in a Glomax 96 microplate luminometer (Promega), resulting in data being presented as relative light units (RLU).

#### RT-PCR and Real-time PCR analysis

RT-PCR and Real-time PCR were carried out as we previously reported (*33*). Briefly, total RNA and DNA were extracted from HepG2 or PHH cells using the RNeasy and DNeasy kits, respectively. On-column DNA digestion was carried out during RNA extraction. For the synthesis of the first strand of cDNA, 1 μg of total RNA after DNase treatment was reverse transcribed using a cDNA synthesis kit. The cDNA samples were then amplified by PCR using 2.5 units of Taq DNA polymerase (Qiagen). Real-time PCR was performed on a CFX96 detection system (Bio-Rad) with SYBR Green reagents (Qiagen). All PCR primers’ sequences are listed below:

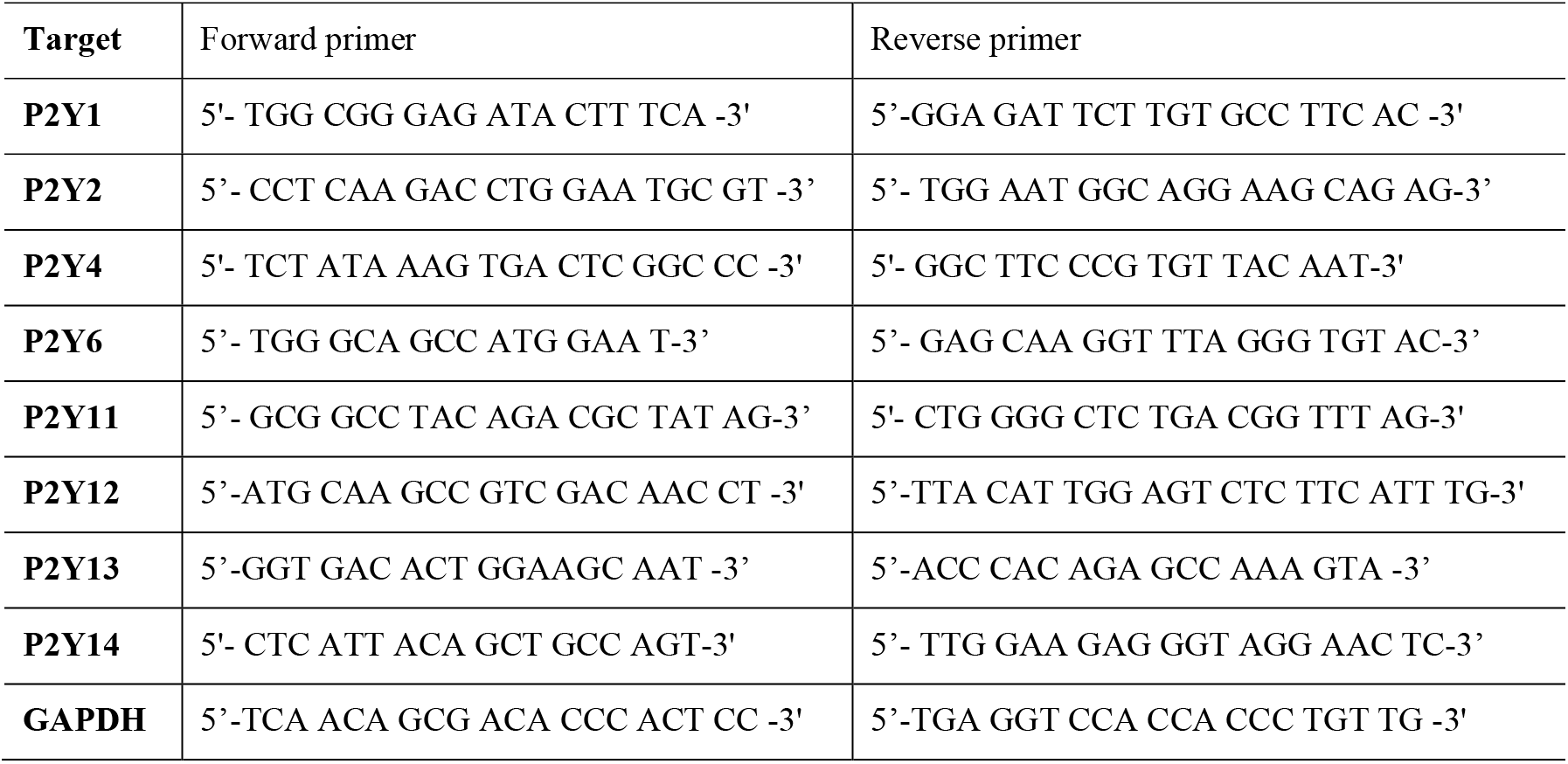

#### Western blotting assays

Western blotting was carried out as we previously reported (*34*). Briefly, after the cells were stimulated for 10 min or indicated times, the cell medium was immediately removed, and the plate was dunked in ice-cold PBS, dried, and quickly lysed with lysis buffer. After collecting lysate in labeled 1.5 mL tubes, the lysate was placed in boiling water for 10 min and immediately placed on ice. Additionally, a 1x solution of SDS running buffer was produced, and 10-well Bio-Rad gel was submerged in running buffer 10 min before removing the comb. Samples were loaded into wells, ran at 70 V for 30 min, and then ran at 110 V for 1 h. While electrophoresis occurred, blotting paper was soaked in transfer buffer, and the PVDF membranes were charged in methanol. Following electrophoresis, proteins were transferred from the gel to a membrane using a semi-dry transfer for 30 min at 20 V. Membranes were collected and washed in TBS-T for 1 min and transferred to a powdered milk solution for 1 h to block the membranes effectively. After blocking, membranes are washed four times for 10 min with TBS-T and placed in primary antibody overnight on a shaker at 4 °C. The individual primary antibodies used were anti-p-AKT (S473/T308), anti-p-ERK1/2, anti-p-p38, and anti-p-JNK (1:1000). Equal protein loading was verified by stripping off the original antibodies and re-probing the membranes with the primary antibody β-actin, GAPDH, total AKT, ERK1/2, p38, or JNK (1:1000). The following morning, membranes in a primary antibody were removed from the 4 °C refrigerators and washed four times for 10 min in TBS-T. They were then placed in a secondary antibody in milk solution for 1 h. Afterward, membranes are washed four times for 10 min in TBS-T and taken to the darkroom for imaging and development on X-ray film. Band densitometry was quantified using ImageJ (NIH), and the ratio-metric calculations were performed in Microsoft Excel and exported to Prism9 for graphing.

#### PIP_2_ supplementation

##### PIP_2_ uptake optimization

HepG2 cells were supplemented with PIP_2_ utilizing a PIP_2_ shuttle assay kit (Echelon Biosciences). To optimize cellular loading of PIP_2_, first, PIP_2_-BODIPY was mixed with a histone H1-TMR carrier at a 1:1 ratio for 15 min at room temperature, vortexing periodically. The fluorophore-bound PIP_2_ complex was diluted to 25 μM in serum-free EMEM. HepG2 cells were then incubated in the complex-containing medium and imaged every 15 min using an EVOS fluorescent imaging system’s GFP and RFP filter to determine the optimal time for BODIPY-PIP_2_ uptake.

##### PIP_2_ treatment for western blotting

HepG2 cells were seeded in 6- or 12- well plates for at least 24 h and cultured in serum-free EMEM 4-6 h before the experiment. HepG2 cells were then treated with 25 μM non-fluorescent PIP_2_-Histone H1 for 30 min, then treated with 1 nM insulin, 100μM nucleotide, or a combination for 10 min. The cells were then lysed and prepared using the aforementioned western blot protocol.

#### In-Cell PIP_3_ ELISA

All reagents and buffers used in PIP_3_ ELISA are listed below. HepG2 cells were seeded in a 96-well plate and starved in a serum-free medium for 16 h before stimulation. Insulin and ATP/UTP were added to the assay wells in the indicated doses incubating for 5 min at 37 °C (the PIP_3_ kinase inhibitor wortmannin was pre-added for 30 min as a positive control). Then, removed the starvation medium and fixed the cells with Fixation Buffer for 15 min at room temperature. Cells were rinsed three times with cold Wash Buffer and treated with Buffer 1 Plus for 45 min on ice in order to block and permeabilize the cells. Next, removed Buffer 1 Plus and added Buffer 2 with Mouse Biotinylated Anti-PI(3,4,5)P_3_ IgM at 1:200 dilution. After 60 min incubation on ice, fixed cells were rinsed once with Buffer 1 and three times with Wash Solution. Dried the plate and added prepared Streptavidin Solution, incubating for 45 min at room temperature with gentle shaking. Cells were then rinsed with Wash Solution four times and incubated with TMB One-Step Substrate Reagent for 30 min at room temperature in the dark with gentle shaking. Finally, Stop Solution was added to each well, and read the absorbance at 450 nm immediately in SpectraMax^®^ iD3 Multi-Mode Microplate Reader.

*Reagents and buffers used in PIP_3_ ELISA*:

| Items | Description | Company |
| --- | --- | --- |
| Mouse Biotinylated Anti-PI(3,4,5)P<sub>3</sub> IgM | 1.0 mg/mL | Echelon Biosciences Inc. |
| Fixation Buffer | 4% formaldehyde and 0.2% glutaraldehyde in PBS | In-house prepared |
| Wash Buffer | 50 mM NH<sub>4</sub>Cl in PBS | In-house prepared |
| Buffer 1 | 20 mM PIPES, pH6.8, 137 mM NaCl, 2.7 mM KCl | In-house prepared |
| Buffer 1 Plus | Buffer 1 supplemented with 5% normal goat serum, 0.5% saponin and 50 mM NH<sub>4</sub>Cl | In-house prepared |
| Buffer 2 | 20 mM PIPES, pH6.8, 137 mM NaCl, 2.7 mM KCl, 5% normal goat serum, 0.1% saponin | In-house prepared |
| Wash Solution | Dilute Wash Concentrate (20x) with DI water | RayBiotech |
| 1x Assay Diluent B | Dilute Assay Diluent B (5x) with DI water | RayBiotech |
| Streptavidin Solution | Dilute HRP-Streptavidin Concentrate (200x) with 1x Assay Diluent B | RayBiotech |
| TMB One-Step Substrate Reagent | 3,3',5,5'-Tetramethylbenzidine (TMB) in buffer solution | RayBiotech |
| Stop Solution | 0.2 M sulfuric acid | RayBiotech |

#### Silencing of P2Y2R by siRNA

To knock down P2Y2R expression, HepG2 cells were transfected with the four-sequence pool (ON-TARGET plus SMART pool L-003688-00-0005, human P2Y2R, NM_002564, Dharmacon) using DharmaFECT-1 Transfection reagent following the manufacturer’s protocol as we previously reported (*32*). Briefly, HepG2 cells were seeded in 6-well plates at 80–90% confluence; the medium was replaced with complete MEM without antibiotics before transfection. DharmaFECT 1 and siRNA products were incubated separately in MEM at room temperature for 5 min. Mixtures were combined, incubated another 20 min, and added to cells at a final concentration of 2 μL/mL DharmaFECT 1 and 25 nM siRNAs. Real-time PCR assays were performed to confirm the decrease of P2Y2R mRNA after 48 h post-transfection. For ATP and UTP stimulation, siRNA and transfection reagent were removed 16 h posttransfection, and a complete culture medium was added. Experiments were performed 72 h post-transfection.

#### Generating HepG2 stable cell line expressing the membrane displayed ATP sensor (pm-iATPSnFR1.1)

The purchased plasmid in bacteria as agar stab for the membrane-displayed ATP sensor for mammalian cells (pm-iATPSnFR1.1) was amplified in LB cultured medium with ampicillin antibiotic at a final concentration of 100 μg/mL. Maxi Plasmid Isolation Kit (Qiagen) was used to isolate and purify the plasmids, of which the sequence was verified by sequencing, and the concentration/purity was measured through NanoDrop2000. The purified plasmids were transfected into HepG2 cells using Optimum MEM medium and Lipofectamine^®^ 3000 Transfection Reagent, after which a stable cell line (HepG2-AS) was produced by G418 selection at 700 ng/mL for two weeks, followed by 400ng/mL for one week until hepatocytes begin growing at a standard rate, then a maintenance dose of G418 at 100ng/mL was used. A functional plasma membrane ATP sensor was verified using the EVOS fluorescent imaging system with the GFP filter, showing the green fluorescent signals at the cellular plasma membrane when the cells were challenged with 30 μM extracellular ATP or 25mM glucose.

#### Observing ATP secretion in HepG2-AS cells

HepG2-AS cells were cultured in EMEM containing 10% FBS and 100ng/mL G418 in a 6- or 12-well plate for 24 h at 70% confluence and then serum starved in antibiotic-free EMEM for 3-5 h before beginning the experiment. NTPDase inhibitor ARL67156 was added to all wells at a concentration of 100μM for 35 min to control for enzymatic ATP degradation across the treatment conditions. Cells were pretreated with 30 μM metformin or a PBS vehicle for 35 min before supplementing 5- or 25 mM D-glucose. HepG2-AS cells were then incubated at 37° C for 1 h and then imaged. Elevated concentrations of extracellular ATP promoted plasma membrane fluorescence of the ATP sensor, which was observed and captured using an EVOS fluorescent imaging system with the GFP filter.

#### Imaging lysosome acidification in HepG2 cells

##### Preparation of pH-sensitive LysoView™ 633 (Biotium)

Lyophilized powder was briefly centrifuged and resuspended in Milli-Q-H2O to generate a 1000X stock solution.

##### Performing lysosome imaging

HepG2 cells were plated in a 6- or 12-well plate after being strained through a 70 μm cell strainer and grown into a monolayer for 24 h. On the day of the experiment, cells were serum-starved for 3-6 h and then pretreated with 30 μM metformin or a PBS vehicle for 35 min before supplementing 5- or 25 mM D-glucose. 30 min following D-glucose supplementation LysoView™ 633 at the final concentration of 1X and CellBrite^®^ Green (Biotium) diluted 1:200 was added to each well and incubated for 30 min at 37° C then imaged using an EVOS fluorescent imaging system with the Cy5 and GFP filters.

#### Intracellular MANT-ATP accumulation

HepG2 cells were plated in a 6- or 12-well plate the day before the experiment and were serum-starved 3-6 h prior to pretreatment with 30 μM metformin or a PBS vehicle for 35 min. The HepG2 cells were then supplemented with 1.5 mM MANT-ATP and incubated for 25 min at 37°C before being imaged using an EVOS fluorescent imaging system with the GFP filter. The captured images were then inverted into black and white using ImageJ.

#### VNUT immunofluorescence assay

Cell immunofluorescence assay was carried out as we previously reported (35). Briefly, HepG2 cells were seeded in 8-chamber glass slides (Nunc), starved for 6 hr, and then the cells were fixed for 15 min in cold methanol. The fixed cells were washed with PBS three times and blocked with 5% horse serum for 1 hr at room temperature. Then the cells were incubated with a rabbit anti-human SLC17A9 (VNUT) antibody (Proteintech) at 1:100 dilution overnight at 4 °C, followed by incubation with an anti-rabbit IgG F(ab) fragment conjugated with AF-488 (Cell Signaling) at 1:1000 dilution for 90 min at room temperature in darkness. For negative controls, cells were incubated with non-immune rabbit IgG in place of the primary antibody or only the AF-488-conjugated secondary antibody. Cells were washed with PBS three times before a mounting medium containing DAPI was added to seal the slides. Images with fluorescent signals in random fields were acquired and captured using an AMG EVOS digital inverted multifunctional microscope.

#### Gluconeogenesis assay

Cell released glucose via gluconeogenesis was quantified using the Amplex Red Glucose Assay Kit (Invitrogen™) as detailed below:

##### Cell culture and medium sample preparation

HepG2 cells were seeded in a 6-well plate, and treatment wells were duplicated to monitor gluconeogenesis and another to control for background glucose production. Hepatocytes grew in a monolayer, reaching approximately 85% confluency. After reaching confluency, starve hepatocytes in serum-free EMEM for 6 h. The base medium utilized was DMEM free of serum, phenol red, glucose, pyruvate, glutamate, and lactate. Both glucose production and background medium required HEPES to be added to the medium at a concentration of 10 mM. The glucose production medium was made by supplementing sodium lactate at a concentration of 10mM. The background medium possessed no gluconeogenic substrates. The P2Y2R antagonist ARC-118929XX and ectonucleotidase inhibitor ARL-67156 were added 45 min before removing the starvation medium. While hepatocytes were starving, a cocktail containing treatment and inhibitors was prepared, after which we removed the starvation medium, washed the cells with PBS pre-warmed to 37 °C, and then added 350 μL pre-made cocktail medium to the appropriate wells to incubate for 3.5 h before collecting medium for glucose quantification.

##### Reagent preparation for Amplex Red glucose assay

Before using, dissolve the Amplex Red vial in 60 μL DMSO – the final reaction volume is 100 μL. Prepare 1X reaction buffer by adding 4 mL of 5X reaction buffer to 16 ml dH2O producing a total volume of 20 mL (~100 assays). Next, a 10U/mL stock solution of horseradish peroxidase (HRP) is prepared (1U is the amount of enzyme that will form 1.0 mg purpurgallin from pyrogallol in 20 sec, pH 6 and 20 °C) by dissolving 1 mL of 1X reaction buffer. Prepare a 100U/mL glucose oxidase stock solution (1U amount of oxidase 1umole of β-d-glucose to glucuronolactone and H_2_O_2_ per 1 min, pH 5.1, 30 °C), dissolving the content of the glucose oxidase vial in 1 mL of 1X reaction buffer. Produce a 400 mM stock solution of glucose (72 mg/mL) and a stock of 20 mM H_2_O_2_ by diluting 3% H_2_O_2_ into the appropriate volume of 1X reaction buffer (22.7 μL of 3% H_2_O_2_ in 977 μL 1X reaction buffer).

##### Performing the Amplex Red glucose assay

Prepare a glucose standard curve (0-200 μM) in 1X reaction buffer with a total volume of 50 μL. The positive control is generated by diluting the 400 mM glucose stock to 200 μM in a 1X reaction buffer. There is also an H_2_O_2_ positive control created by diluting 20 mM H_2_O_2_ to 10 μM in 1X reaction buffer. The negative control is 1X reaction buffer without H_2_O_2_. In vitro samples require no dilution prior to reaction. Transfer 50 μL of sample supernatant into their respective wells, ensuring to do a minimum of triplicates. Next, combine 50 μL 100 μM Amplex Red, 100 μL 0.2 U/mL HRP, 100 μL 2U/mL glucose oxidase, and 4.75 mL 1X reaction buffer. Add 50 μL of the Amplex Red cocktail into all wells and allow to incubate protected from light for 30 min at room temperature. Read plate (λab: 571 nm, λem: 585 nm).

#### Glycogen measurement

Cellular glycogen content was evaluated using the Glycogen Assay Kit (Cayman Chemical) according to the manufacturer’s instructions. Briefly, HepG2 cells were seeded in the growth medium in 6-well plates reaching about 80% confluency, and then the cells were serum-starved for 6 h. Before lysing cells, immediately dunk the plate in ice-cold PBS, knock off residual PBS, and carefully snap freeze the 6-well plate in liquid nitrogen. Add 200 μL of premixed 1% Triton-X lysis buffer, vigorously scrape each well, and mechanically lyse cells using a 20-gauge syringe at 4 °C. Then, transfer the samples into a 1.5ml Eppendorf tube, centrifuge at 15,000 rpm for 10 min at 4 °C, and collect the supernatant. After that, 10 μL of cell lysates were added to every well of the 96 wells plate, and then 50 μL of hydrolysis buffer with amyloglucosidase was added to convert glycogen to β-D-glucose. Glucose oxidase was then added to convert β-D-glucose to hydrogen peroxide. Horseradish peroxidase was added, and H_2_O_2_, in the presence of it, reacted with 10-acetyl-3,7-dihydroxyphenoxazine to produce resorufin, whose fluorescence was measured at Ex/Em of 535/590nm. Data were extrapolated based on the standard curve constructed by the kit-provided glycogen standards.

#### Glucose uptake assay

Samples were processed as detailed in the manual of the 2-NBDG Glucose Uptake Assay Kit (Abcam). Briefly, hepatocytes were plated in EMEM medium in 6-well plates reaching about 80% confluency, and then the cells were serum-starved for 6 h. Following starvation, replace the serum-free medium with pre-warmed glucose-free DMEM, and hepatocytes were treated with ATP or UTP for 30 min in total, during which the final 10 min of the cells were incubated with 2-NBDG at the final concentration of 200 μg/mL. After NBDG was taken up by the cells, fluorescence was determined immediately with an excitation wavelength set to 485 nm and an emission wavelength set to 535 nm.

#### Extracellular glucose quantification

The extracellular level of glucose was determined using the Glucose Assay Kit (Abcam). Briefly, HepG2 cells were seeded in a 6-well plate, reaching approximately 85% confluency, after which the cells were starved in serum-free EMEM for 6 h. A standard curve for glucose was produced using the range 0-300 mg/dL. 5 μL of sample supernatants and standards were transferred to 1.5 mL tubes. Add 500 μL of acetic acid-based reagent, close the lids very tightly, and vortex. Notably, due to the acetic acid present in the reagents, this experiment was predominately performed in fume hood—transfer tubes to either a heat block at 100 °C or a boiling water bath for 8 min. Immediately, transfer the tubes to a cool water bath or ice for 4 min. After cooling, transfer 200 μL of the sample, standard, and pure water (blank) to a 96-well plate—measure optical density (OD) at 630 nm. Data were extrapolated based on the standard curve constructed by the kit-provided glucose standards.

#### Animal procedures

Mice homozygous for inactivation of the *P2Y2R* gene (P2Y2R KO, Jackson Laboratory) were back-crossed more than 10 generations into the C57BL/6 background. At 4 weeks, age-matched male mice (P2Y2R-KO versus P2Y2R-WT) were fed either a standard chow diet (ENVIGO) or a high-fat diet (Fat: 42% of total Kcal; Sucrose: 34.5% of total weight; Saturated fatty acid: 66% of all fat; ENVIGO) for another 16 weeks before being used for experiments. All mice were maintained in a facility free of well-defined pathogens under the supervision of the Biological Resource Unit at Auburn University. All animal protocols were approved by Auburn Institutional Animal Care and Use Committee, and the investigation conforms to the Guide for the Care and Use of Laboratory Animals published by the United States National Institutes of Health.

#### Glucose tolerance test

Mice were fasted for 12 h with free access to drinking water. Each mouse then received an intraperitoneal injection of filter-sterilized D-glucose (1.5g/kg body weight) in normal saline (0.9% NaCl). Blood glucose levels were measured using a glucose meter (True Metrix Go^®^ glucometer, TRIVIDIA HEALTH) from tail blood at 0, 15, 30, 60, 90, and 120 min after intraperitoneal injection of glucose solution.

#### Insulin tolerance test

After 6 h fasting with free access to drinking water, mice were given intraperitoneal injections of human insulin (Humulin^®^ U-100, Eli Lilly) at a dose of 0.75 IU/kg body weight using a sterile insulin syringe. Blood glucose levels were determined using a glucose meter (True Metrix Go^®^ glucometer, TRIVIDIA HEALTH) from tail blood at 0, 15, 30, 60, and 90 min after insulin injection.

#### Immunohistochemistry

After euthanizing the mice, the liver was isolated and fixed in 10% neutral buffered formalin. The fixed liver was then refrigerated at 4 °C. After 48 h refrigeration, the liver was transferred to a 30% sucrose solution in PBS, allowing the liver to soak for 48 h in the sucrose solution before sectioning. The liver tissue was then dried and integrated into a mold of frozen section compound (FSC, Surgipath). Liver sections were collected at 10 μm and attached to charged slides (Leica). Slides were then left at room temperature for 1 h before storing at −20 °C. The liver tissue slides were stained using the Super Plus TM High Sensitive and Rapid Immunohistochemical Kit (Elabscience) according to the kit’s instructions. Briefly, mouse liver tissue slides were placed in an Antigen Retrieval working solution for 30 min to repair the antigen. Slides were blocked with peroxidase blocking buffer at room temperature for 15 min to eliminate endogenous peroxidase activity, then washed with PBS three times for 2 min. The primary anti-P2Y2R(ABclonal) or anti-VNUT antibody (Proteintech) with a 1:400 dilution using antibody dilution buffer was added to the slides and incubated at 37°C for 1 h, then washed with PBS three times for 2 min. A vehicle incubation without a primary antibody was used as a control. A drop of Poly-peroxidase-anti-Rabbit/Mouse IgG was added to the slides and incubated at 37°C for 30 min, then washed with PBS three times for 2 min. A drop of coloration solution was added to the slides, and the color tan or brownish yellow appeared. Slides were washed with deionized water at the same time to terminate the chromogenic reaction, after which the sections were counter-stained with Hematoxylin and cover-slipped with VectaMount^®^ AQ Aqueous Mounting Medium (Vector^®^ Laboratories). All images were then acquired and captured with SWIFTCAM Microscope Digital Camera SC1630.

#### Data analysis

All data were analyzed by Prism 9 (GraphPad Software Inc). Data are expressed as the mean ± SEM. The means of two groups were compared using Student’s t-test (unpaired, two-tailed), and one-way analysis of variance was used for comparison of more than two groups with P < 0.05 considered to be statistically significant. Unless otherwise indicated, all experiments were repeated at least three times.

**Fig. S1.**
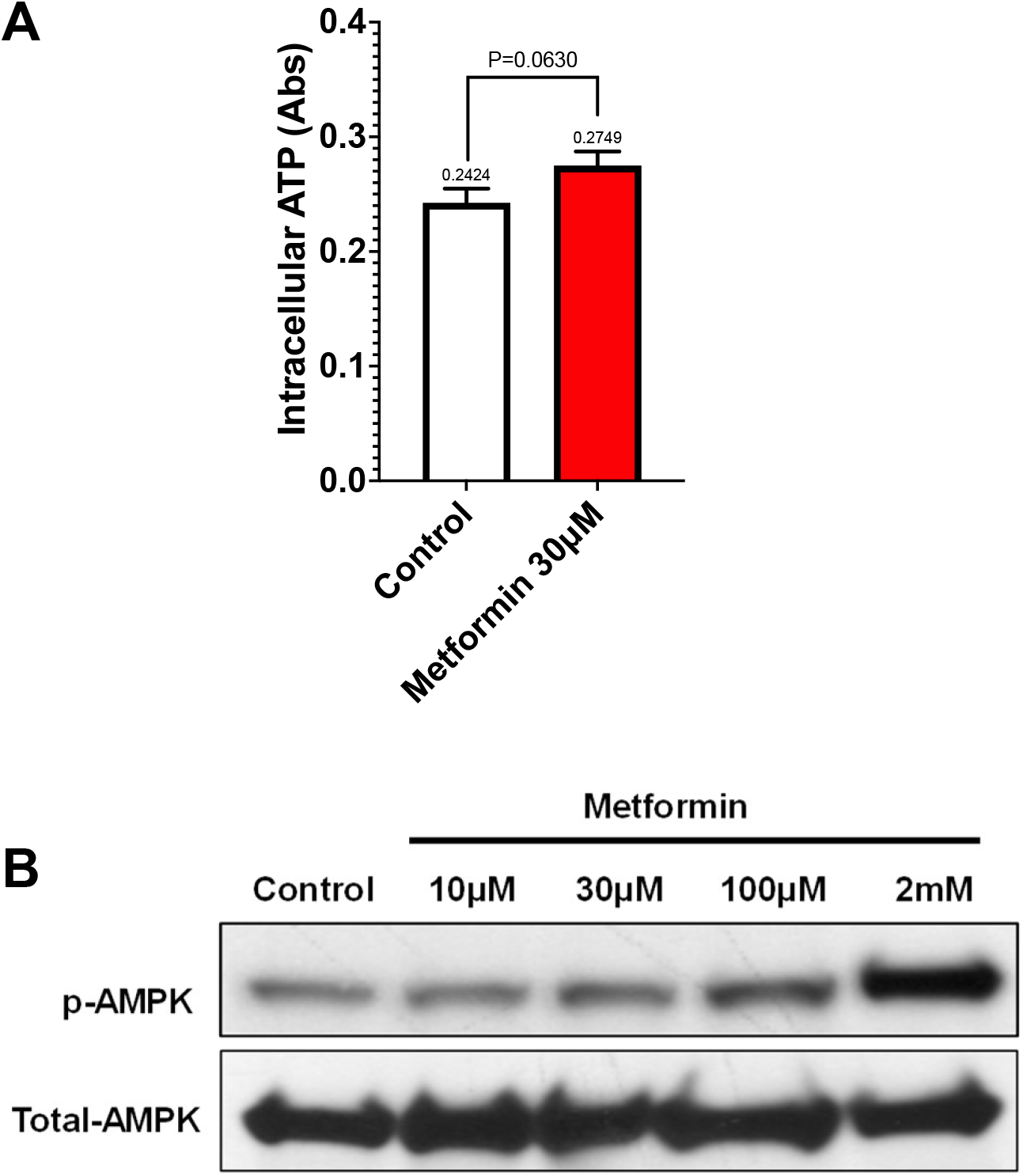
Effect of metformin on intracellular ATP level and AMPK activation in HepG2 cells. (**A**) Intracellular ATP content was assessed following metformin treatment for 2hr at 30μM by an ATP assay kit from Abcam. Cell lysates from the sample wells underwent a deproteination sample preparation step. Measurement of ATP was executed by combining 50 μL assay reagent with 50 μL lysate extracted from each well in a black/clear bottom 96-well plate. Incubated in darkness at room temperature for 30 min. Data was collected from a 96-well plate reader at 570 nm and reported in optical density absorbances (Abs). n=3, each in triplicates. (**B**) HepG2 cells were serum-starved for 6hr, after which the cells were stimulated with the indicated concentrations of metformin for 24hr, followed by a standard Western blotting assay to detect the phosphorylated AMPK (Thr172) and total AMPK. Shown are representative data from three independent experiments.

**Fig. S2.**
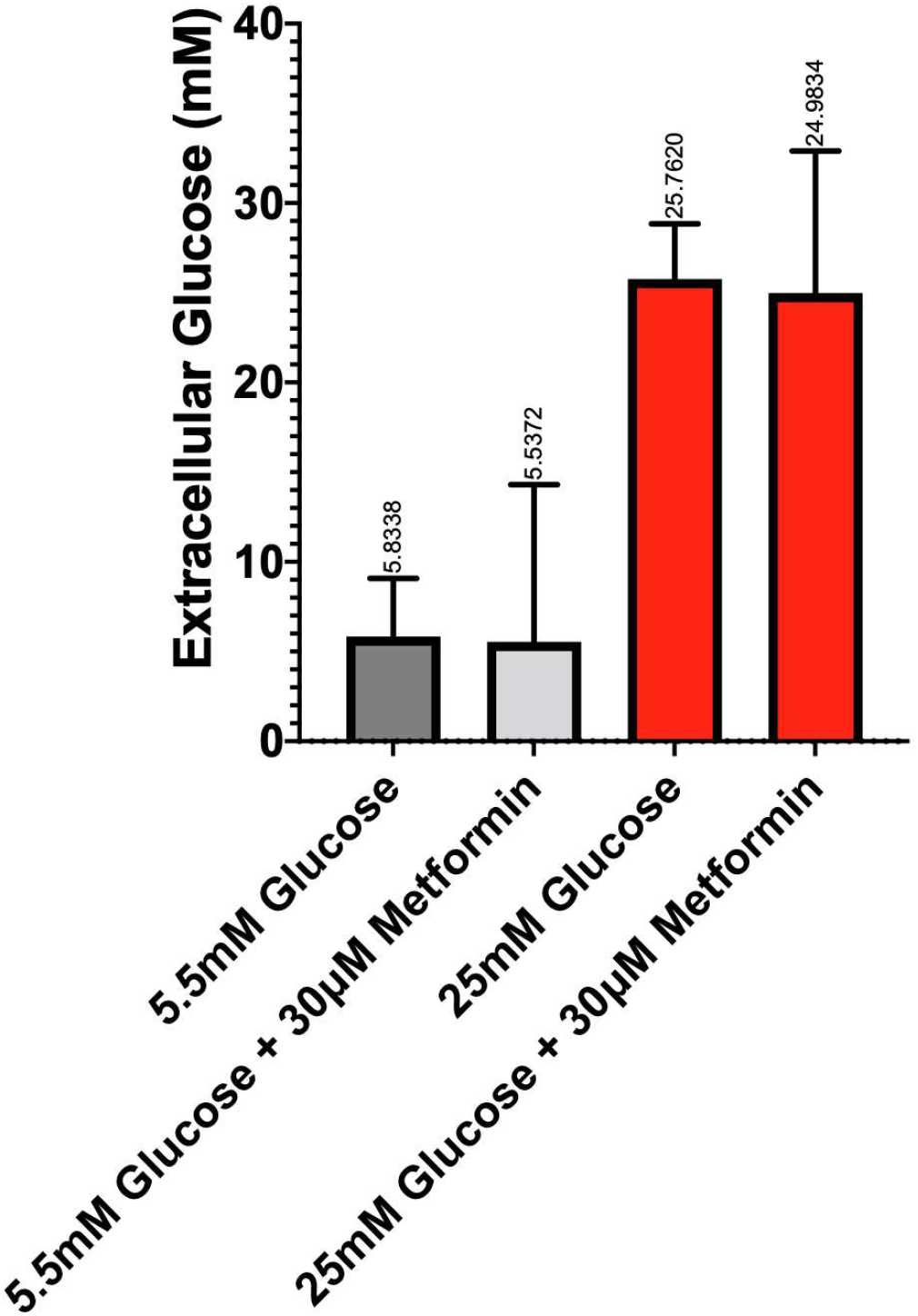
Effect of metformin on extracellular glucose concentration after high glucose challenge of HepG2 cells. Extracellular glucose concentrations were monitored under the indicated assay conditions of 30μM metformin, 5.5mM glucose, and 25mM glucose using a Glucose Assay Kit (ab272532). The cells were pretreated with metformin for 45 min and then challenged with 25 mM glucose for 2hr, after which 5 μL cell culture medium was mixed with the assay reagent to generate the color reaction, of which the absorbance at 630nm is directly proportional to glucose concentration in the sample. Data shown are means ± SEM of three independent experiments and no significant statistical difference was observed between the absence and presence of 30μM metformin.

**Fig. S3.**
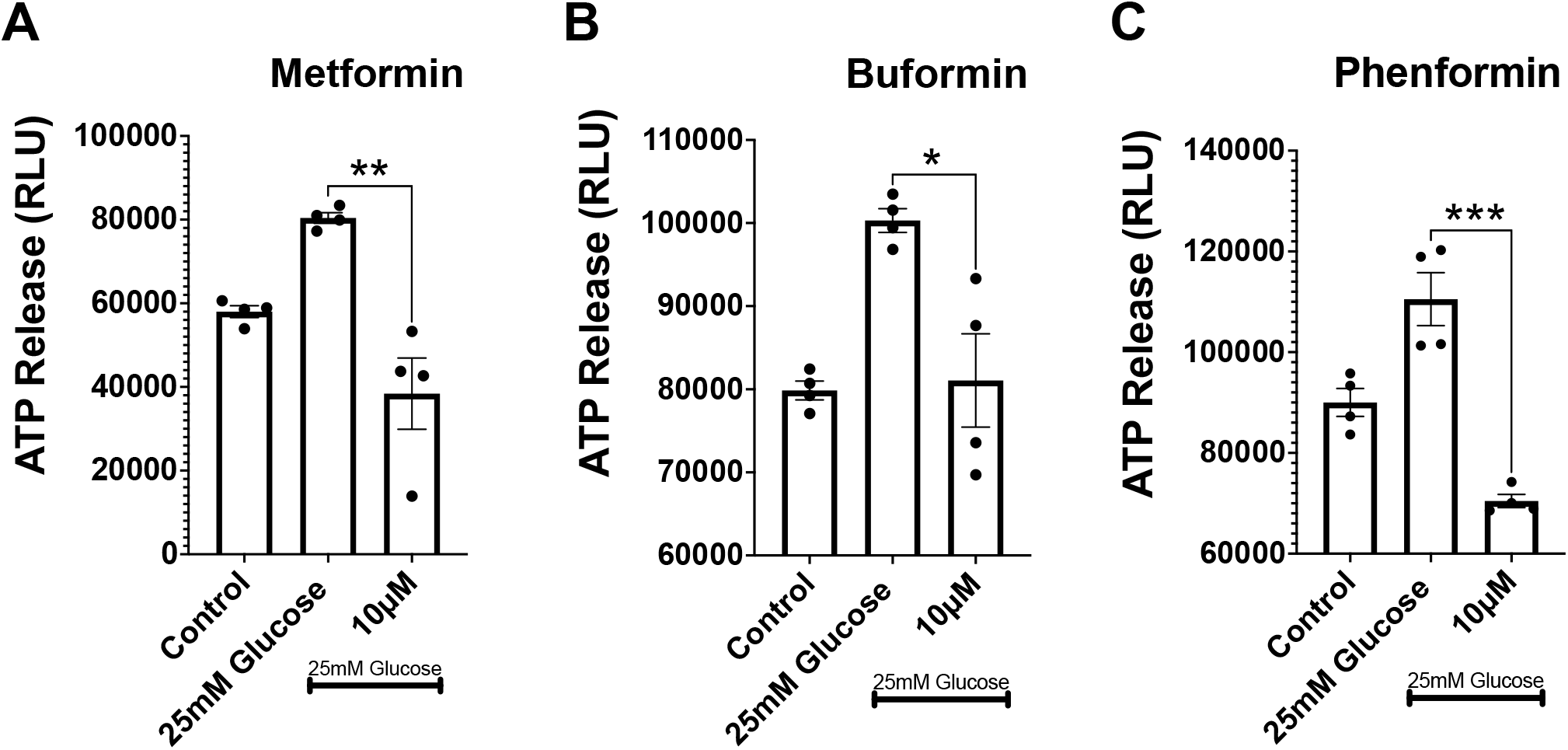
Effect of antidiabetic biguanides on high glucose-stimulated ATP secretion in primary human hepatocytes. Cultured human primary hepatocytes were serum-starved for 6 hr, then pretreated with the indicated concentration of metformin (**A**), buformin (**B**), or phenformin (**C**) for 35 min, followed by high glucose challenge for 1.5 hr, after which the cell culture medium were taken for ATP quantification by luciferin-luciferase assay. Data are means ± SEM of four independent experiments. *P<0.05, **P<0.01, ***P<0.001.

**Fig. S4.**
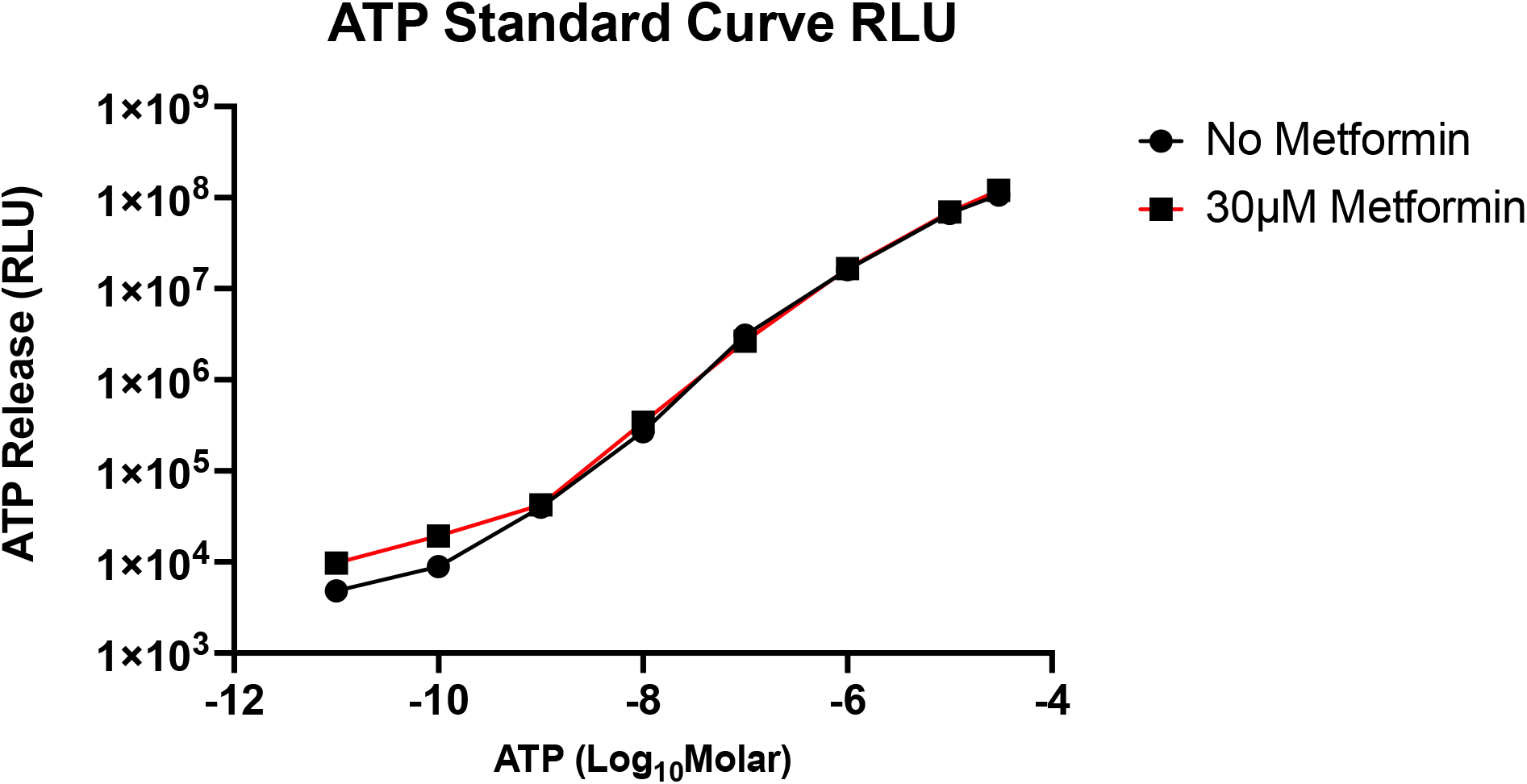
Effect of metformin on the ATP standard curves measured by luciferin-luciferase assays. Measurement of ATP standard curves was executed utilizing the ENLIGHTEN ATP Assay System Bioluminescence Detection Kit by combining 100 μL assay reagent with 100 μL cell culture medium containing all the mentioned components but without HepG2 cells in a black/clear bottom 96-well plate. Quantification was accomplished in a Glomax 96 microplate luminometer, resulting in data presented as relative light units (RLU). Data shown are means ± SEM of three independent experiments and no significant statistical difference was observed between the absence and presence of 30μM metformin.

**Fig. S5.**
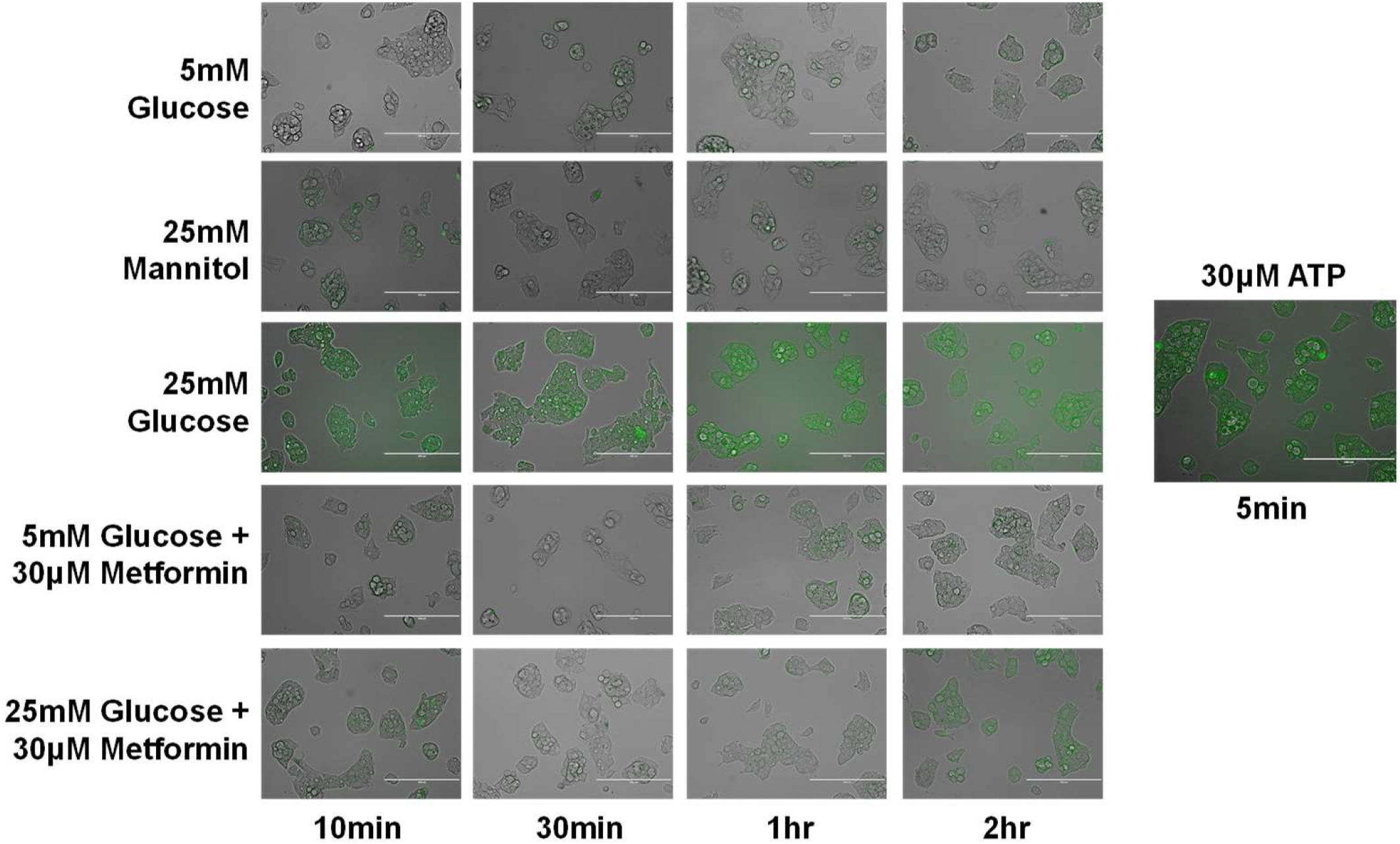
Effect of metformin on high glucose-stimulated ATP secretion in HepG2 hepatocytes. Qualitative analysis of ATP secretion in HepG2 cells transfected with a Bioengineered Cell Surface ATP Sensor (iATPSnFR) (green) exposed with the indicated times and concentrations of glucose and metformin. 30μM ATP was used as a positive control for the transfected ATP sensor. The data shown are representative of three independent experiments. Scale bar: 200μm.

**Fig. S6.**
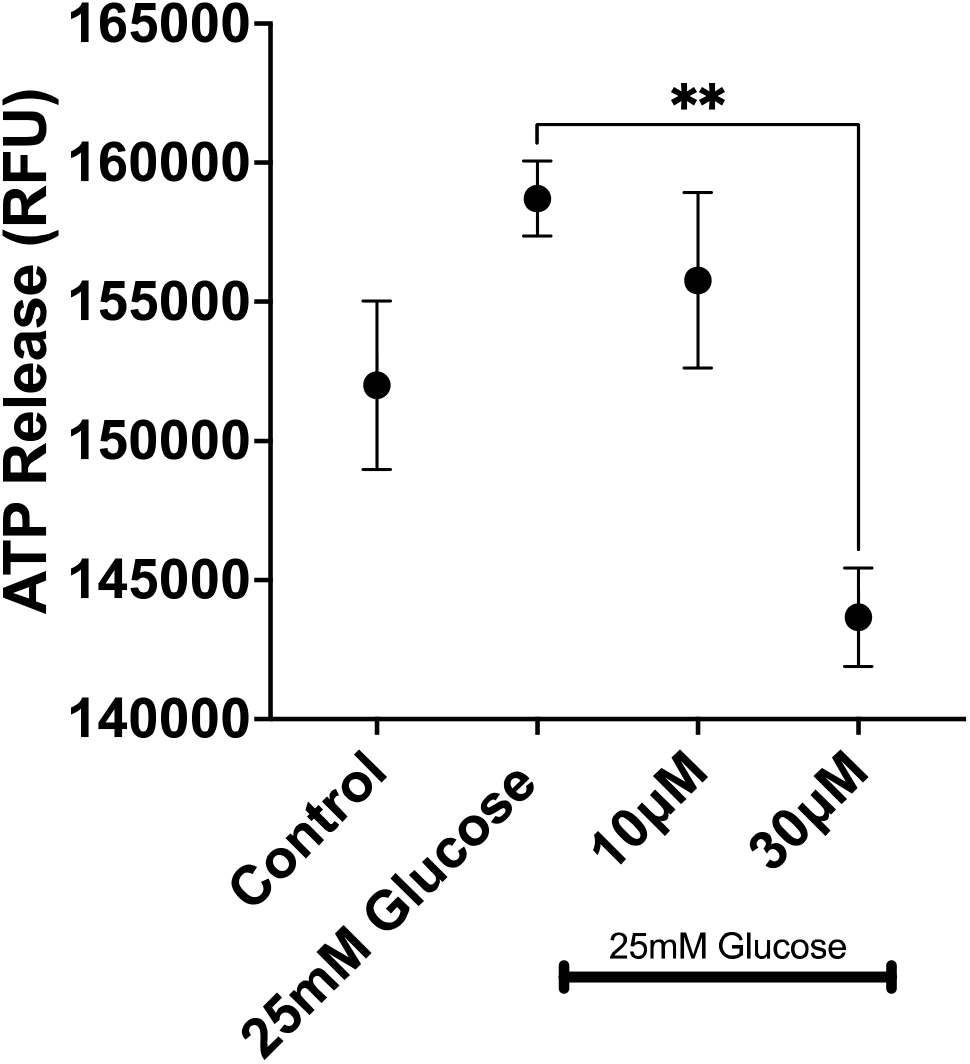
Metformin inhibition of GSAS in HepG2 cells measured by fluorometry. ATP secretion from HepG2 cells was quantified utilizing a colorimetric/fluorometric ATP Assay Kit from Abcam. Hepatocytes were cultured in a 6-well plate 24 h prior to experimentation and starved for 4 h before the start of the experiment. NTPDase inhibitor ARL67156 was added to every well at a concentration of 100μM for 30 min before starting the experiment. Metformin was pretreated for 35 min at the indicated concentrations before challenging the cells with 25 mM D-glucose. Measurement of ATP was executed by combining 50 μL assay reagent with 50 μL medium extracted from each well in a black/clear bottom 96-well plate. Incubated in darkness at room temperature for 30 min. Data was collected from a 96-well plate reader using Ex/Em = 535/587nm and reported in relative fluorescent units (RFU). Data are means ± SEM of three independent experiments. **P<0.01

**Fig. S7.**
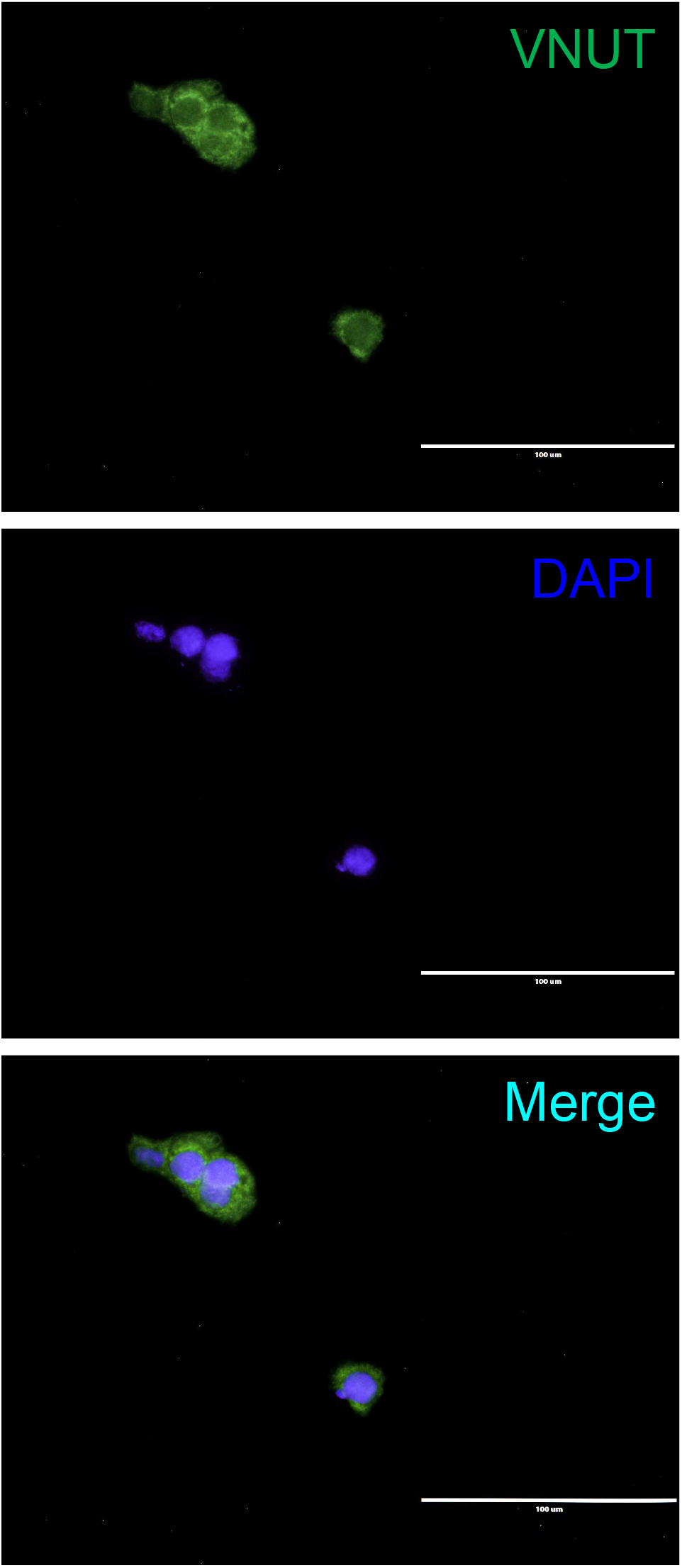
Evidence of VNUT expression in HepG2 hepatocytes. Immunofluorescent validation of VNUT (green) expression relative to nuclear DAPI (blue) in HepG2 hepatocytes. Cultured HepG2 cells were serum-starved for 6 hr, fixed in pre-cooled methanol, followed by a standard immunofluorescent assay using a rabbit anti-human SLC17A9 (VNUT) antibody at 1:100 dilution and an anti-rabbit IgG F(ab) fragment conjugated with AF-488 at 1:1000 dilution. The data shown are representative of three independent experiments. Scale bar: 100μm.

**Fig. S8.**
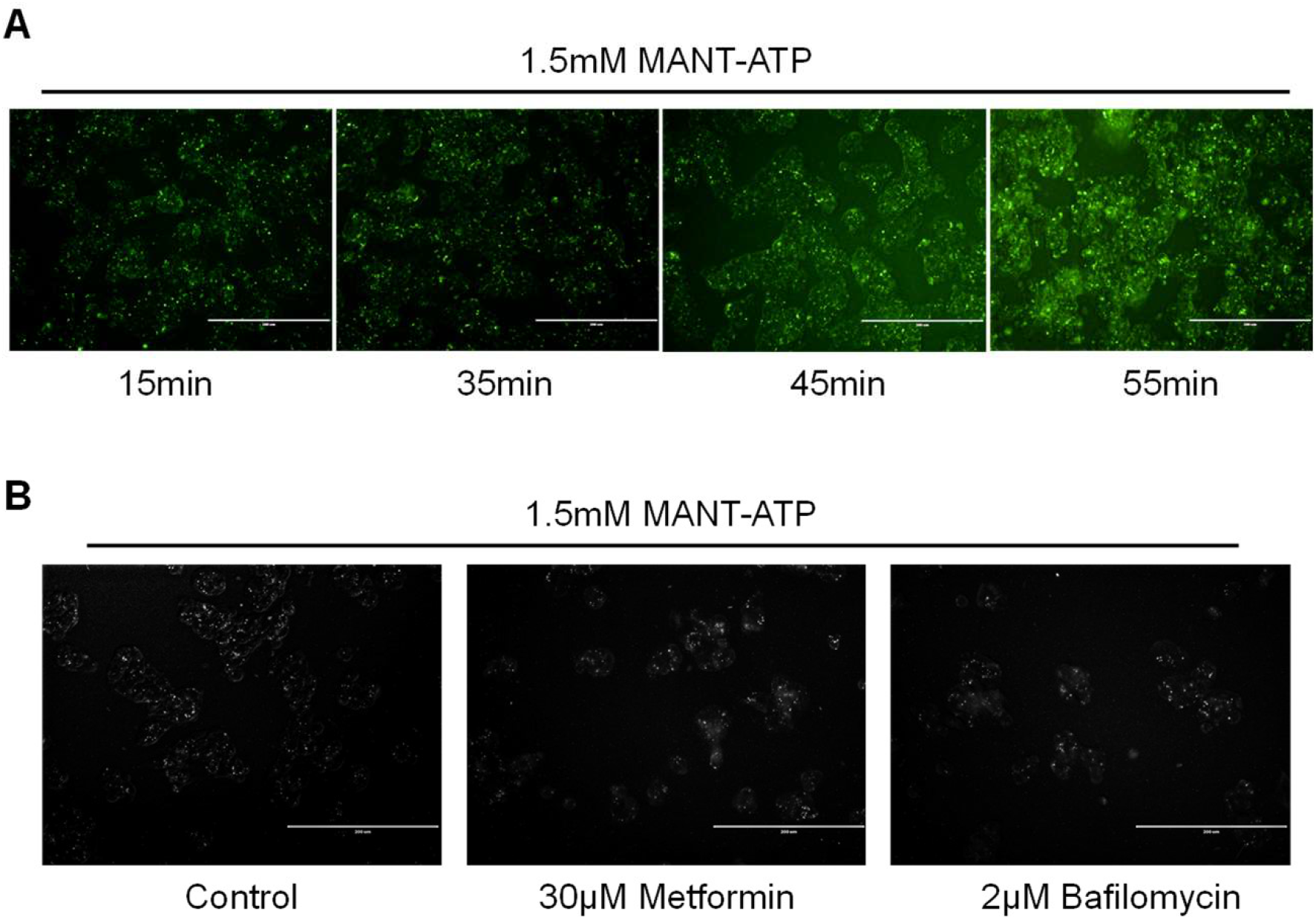
Effect of metformin on MANT-ATP uptake in HepG2 hepatocytes. (**A**) Culture HepG2 cells were serum-starved for 6 hr, then 1.5 mM MANT-ATP was added, after which green fluorescent signals representative of MANT-ATP uptake were detected at the indicated time points. (**B**) Cultured HepG2 cells were serum-starved for 6 hr, then pretreated with the indicated concentration of metformin or bafilomycin or vehicle control for 35 min, followed by adding 1.5mM MANT-ATP and incubated for another 30min at 37°C before acquiring the images. For clarity, the original green images were transformed to black/white format by Image J2. The data shown are representative of three independent experiments. Scale bar: 200μm.

**Fig. S9.**
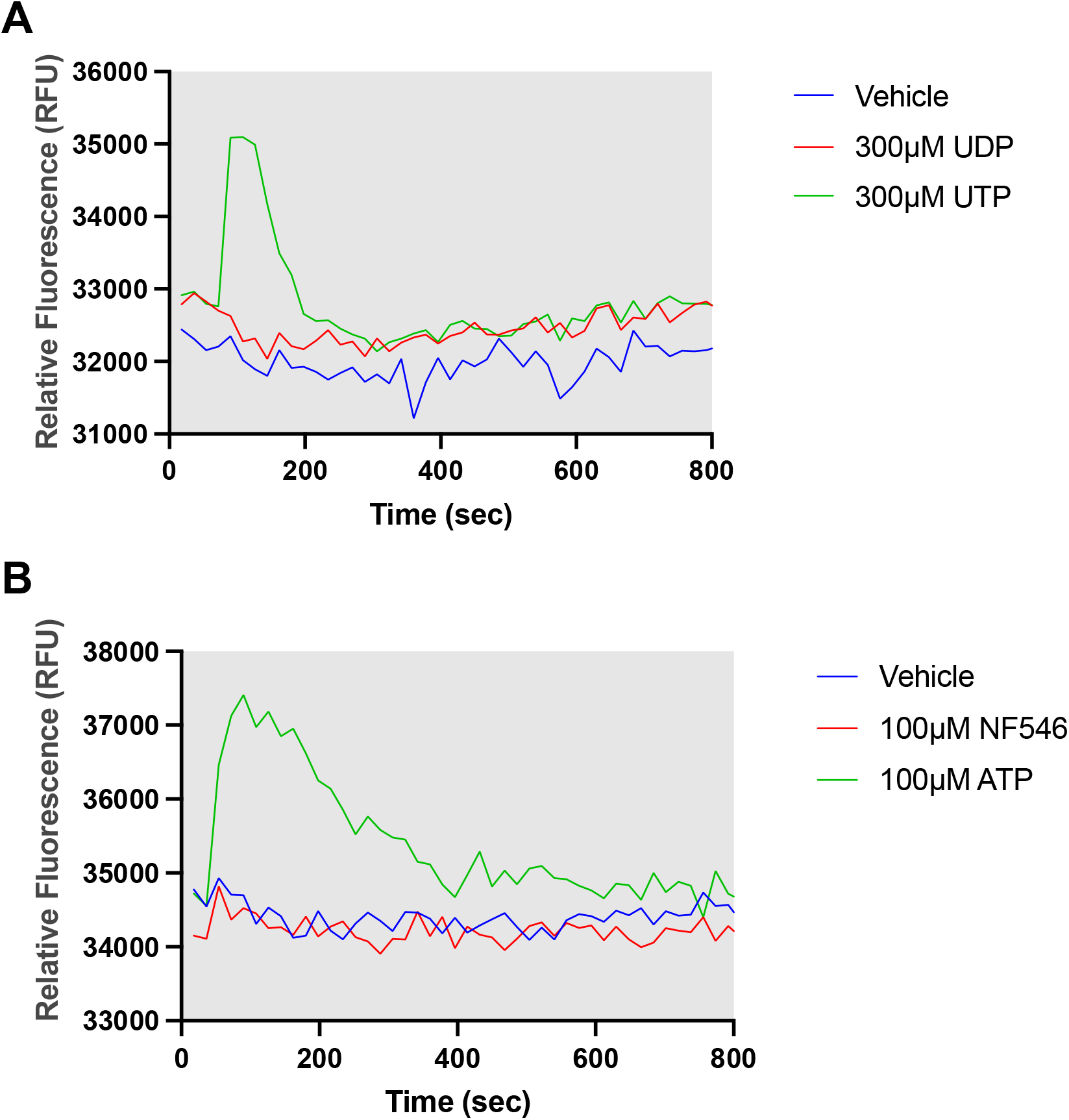
Evidence of no functional P2Y6 and P2Y11 receptors in HepG2 cells. HepG2 cells were stimulated with P2Y6R (**A**) and P2Y11R (**B**) agonists UDP and NF546, respectively, at the indicated concentrations, and intracellular [Ca^2+^]_i_ mobilization was monitored fluorometrically relative to P2Y2R agonist ATP and UTP using the FluoForte™ Calcium Assay kit. The time-response curves of intracellular [Ca^2+^]_i_ signal were recorded via real-time monitoring of fluorescence intensity (λex: 490 nm, λem: 525 nm) in a Fluorometric Microplate Reader. Shown are representative tracings from three independent experiments.

**Fig. S10.**
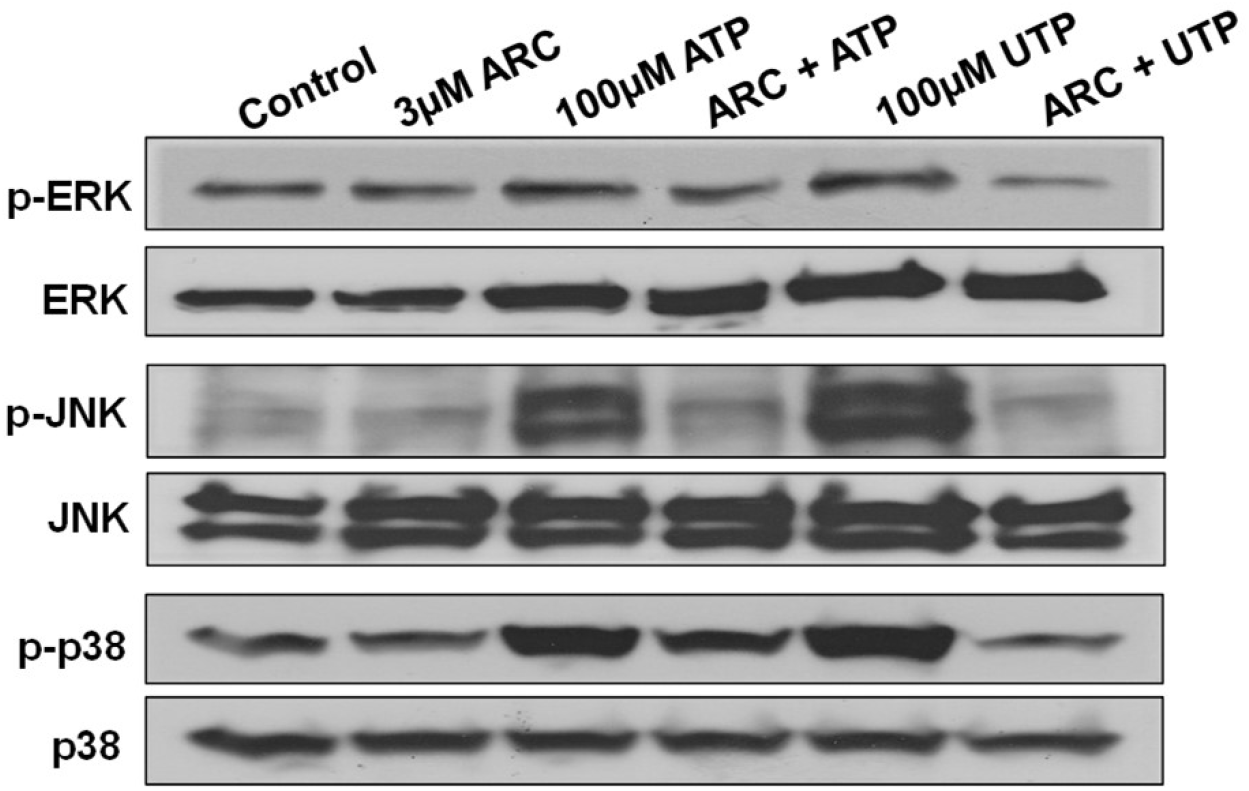
Effect of P2Y2R-selective antagonist ARC-118925XX on nucleotide-induced MAPK signaling in HepG2 cells. HepG2 cells were serum-starved overnight, after which the cells were pretreated with the P2Y2R-selective antagonist ARC-118925XX at 3μM for 45 min, followed by ATP or UTP stimulation at the indicated concentration for 10 min, followed by a standard Western blotting assay to detect the phosphorylated MAPK and total MAPK. Shown are representative data from three independent experiments.

**Fig. S11.**
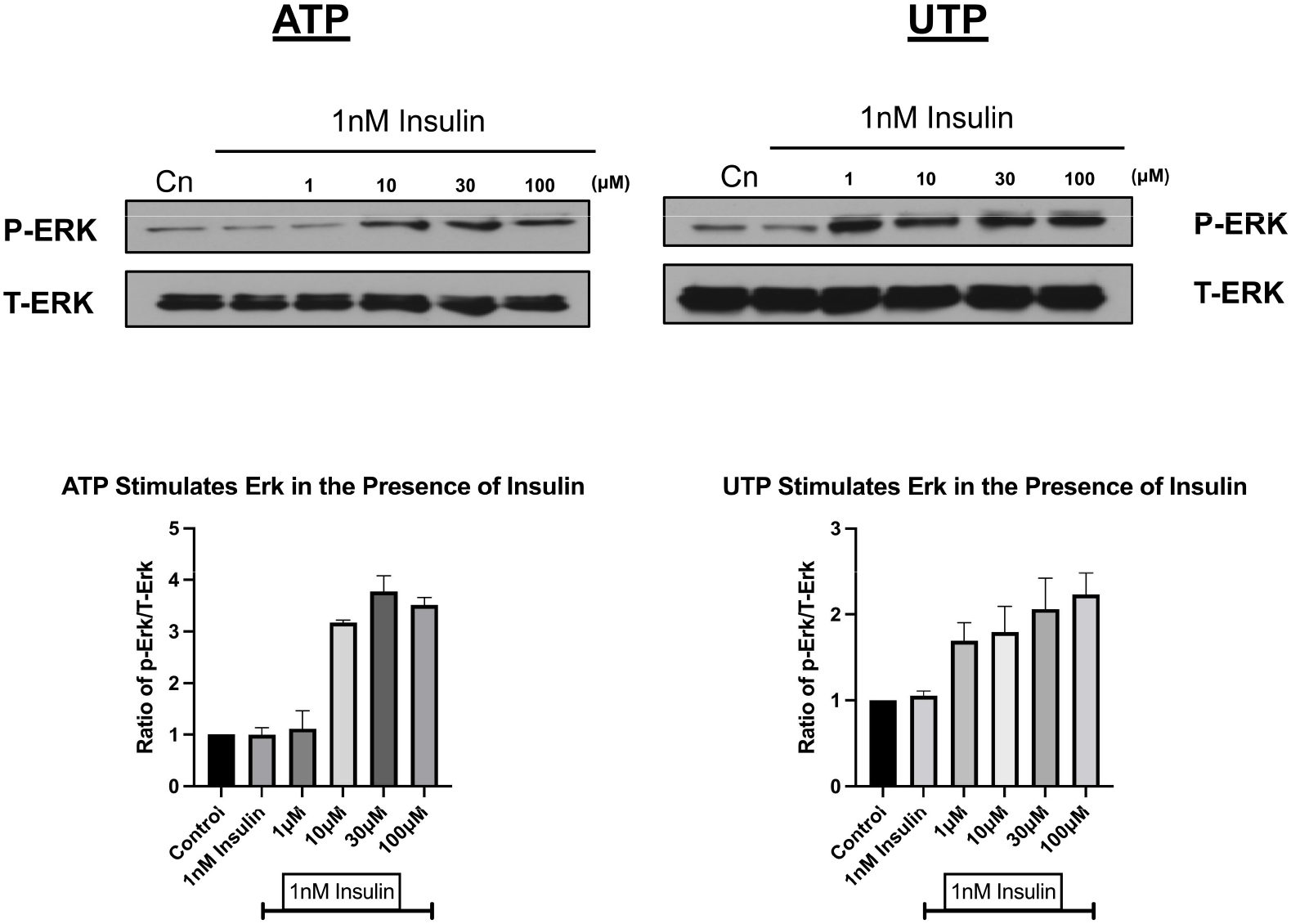
Effect of insulin on P2Y2R-mediated ERK1/2 activation in HepG2 cells. HepG2 cells were serum-starved overnight, after which the cells were stimulated by the indicated concentrations of ATP or UTP together with 1 nM insulin for 10 min, followed by a standard Western blotting assay to detect the phosphorylated ERK1/2 and total ERK1/2. Shown on top are representative data from three independent experiments and the mean densitometric analysis of three separate experiments are normalized to respective total ERK1/2 loading controls and the summarized data were shown in the bottom graph.

**Fig. S12.**
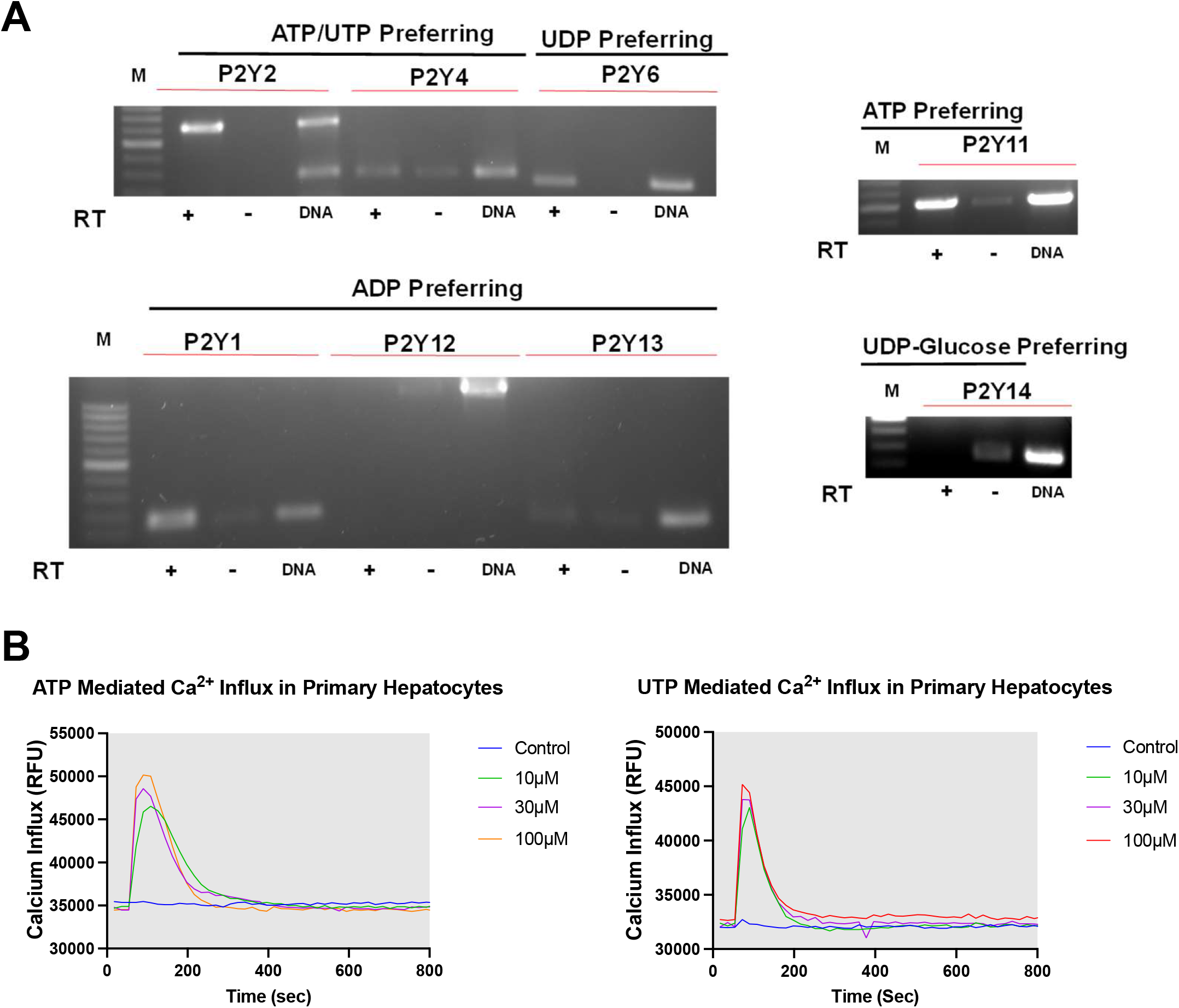
Evidence of P2Y2R expression in primary human hepatocytes. (**A**) RT-PCR analysis of mRNA expression profile of the eight known P2Y receptor subtypes in primary human hepatocytes. (**B**) Human primary hepatocytes were stimulated with P2Y2R agonists ATP and UTP at the indicated concentrations, and [Ca^2+^]_i_ mobilization was monitored fluorometrically. Shown are representative data from three independent experiments.

**Fig. S13.**
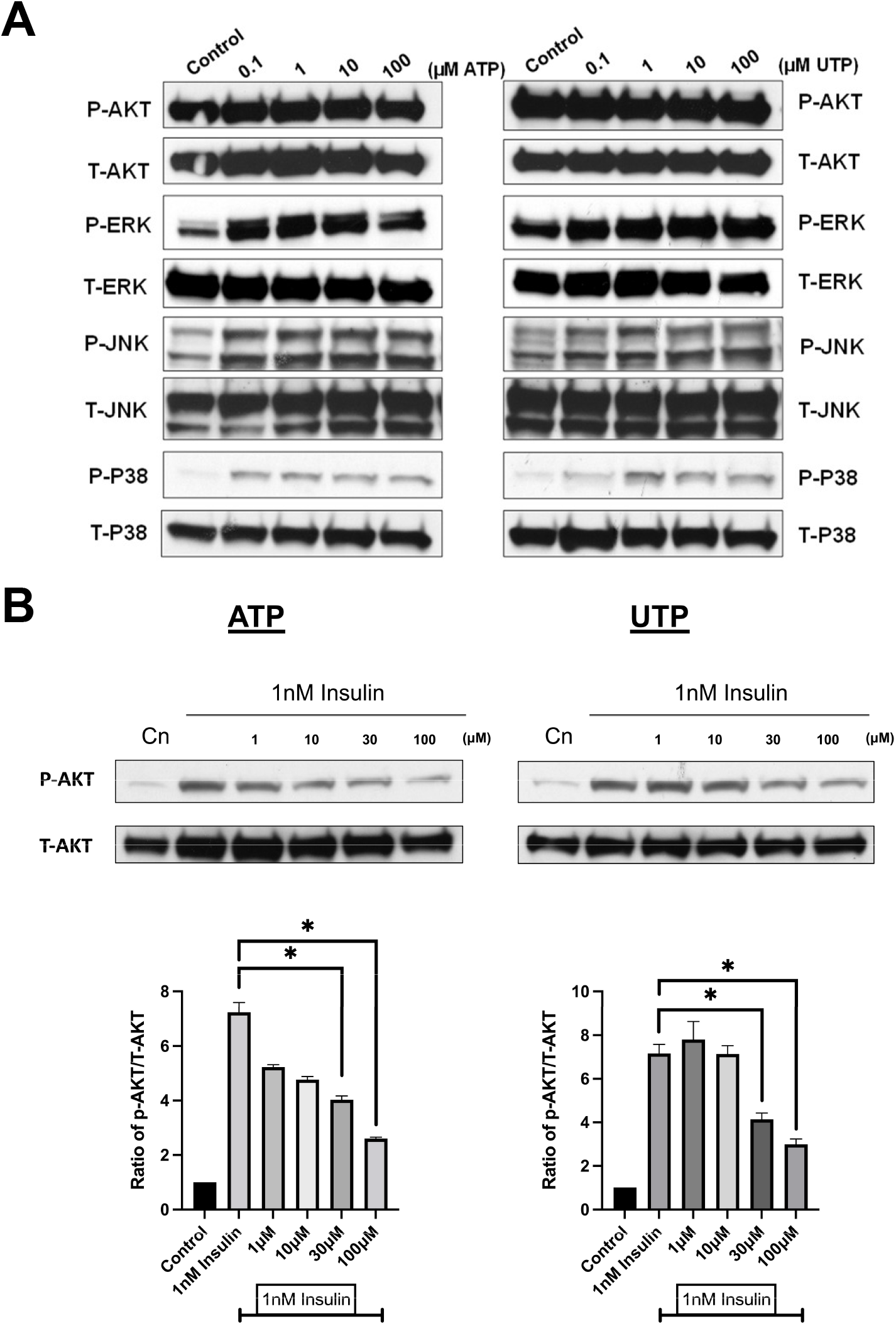
P2Y2R activation of MAPK pathways and its impact on insulin-induced AKT signaling in primary human hepatocytes. (**A**) Primary human hepatocytes were serum-starved for five hours, after which the cells were stimulated by ATP or UTP at the indicated concentrations for 10 min, followed by a standard Western blotting assay to detect the phosphorylated AKT or MAPK and their respective total kinase proteins. Shown are representative data from three independent experiments. (**B**) Primary human hepatocytes were serum-starved for five hours, after which the cells were stimulated by the indicated concentrations of ATP or UTP together with insulin for 10 min, followed by a standard Western blotting assay to detect the phosphorylated AKT and total AKT. Shown on top are representative data from three independent experiments and the mean densitometric analysis of three separate experiments is normalized to respective total AKT loading controls and the summarized data were shown in the bottom graph. * p<0.05.

**Fig. S14.**
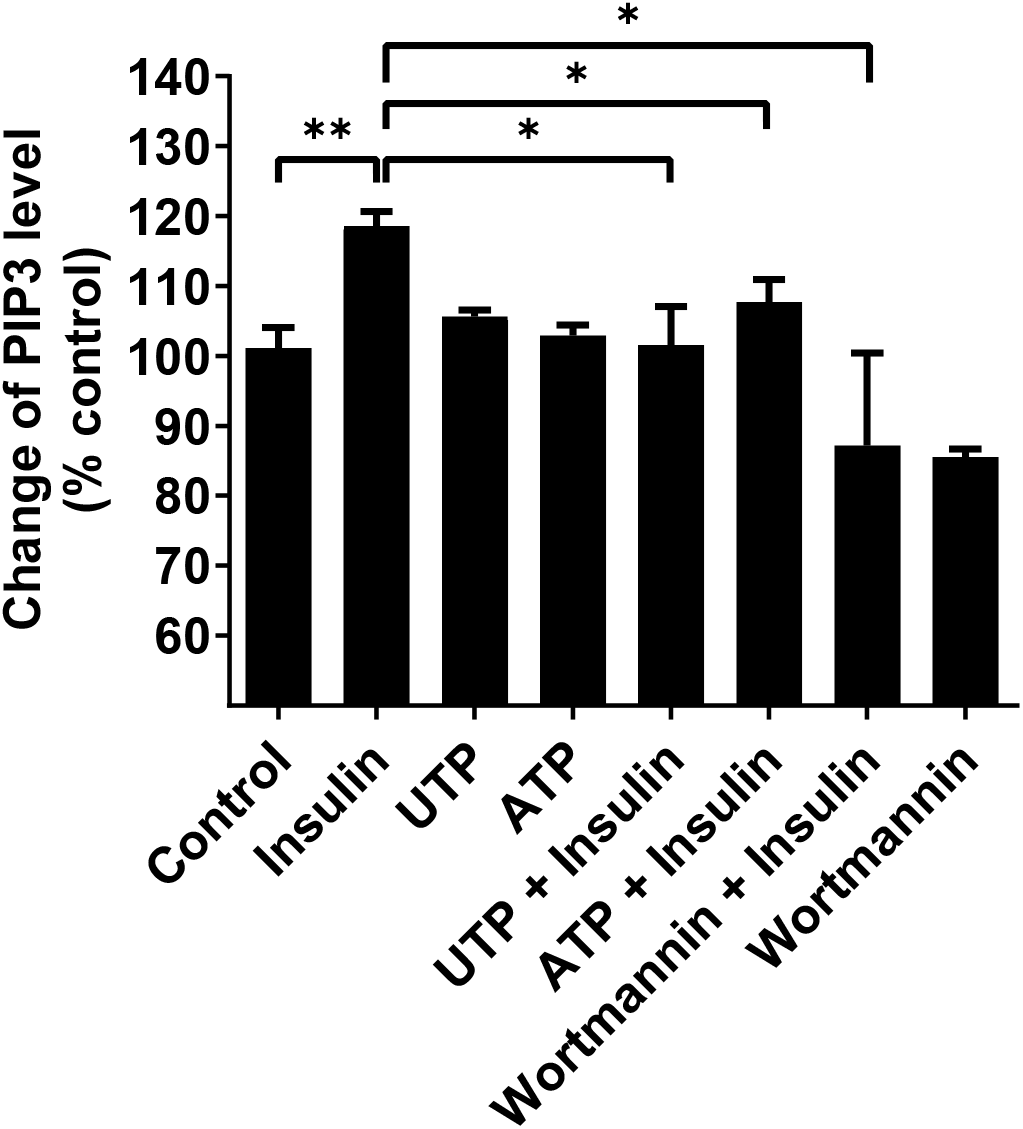
Effect of P2Y2R activation on insulin-induced PIP_3_ accumulation in HepG2 cells. Cells were seeded in a 96-well plate and starved for 16 h before stimulation by 5nM insulin with or without 100μM ATP or UTP for 5 min, after which the cells were fixed and membrane PIP_3_ levels were quantified by ELISA using a biotinylated anti-PIP_3_ IgM at 1:200 dilution, followed by HRP-streptavidin and TMB substrate detection. The control group’s absorbance value at 450 nm reading was considered 100%. The PIP_3_ kinase inhibitor wortmannin was pre-added for 30 min as a positive control. Data are means ± SEM of three independent experiments. *P<0.05, **P<0.01.

**Fig. S15.**
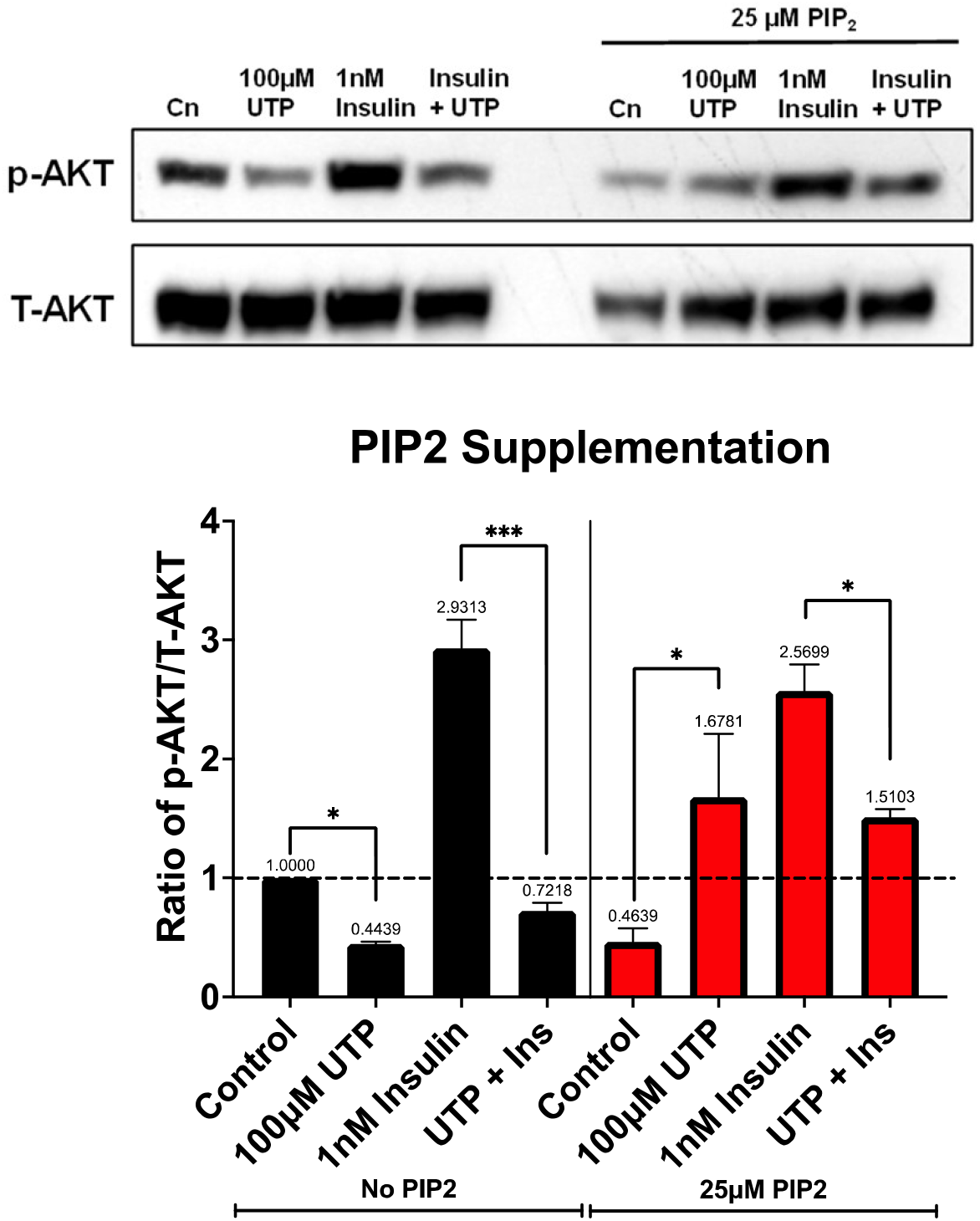
Effect of PIP_2_ supplementation on P2Y2R-mediated suppression of insulin-AKT signaling in HepG2 cells. HepG2 cells were serum-starved overnight, treated with 25 μM PIP_2_ or the carrier control vehicle for 30min, after which the cells were stimulated by the indicated concentration of UTP together with or without 1 nM insulin for 10 min, followed by a standard Western blotting assay to detect the phosphorylated AKT (S473) and total AKT. Shown on top are representative data from three independent experiments and the mean densitometric analysis of three separate experiments are normalized to respective total AKT loading controls and the summarized data were shown in the bottom graph. * P < 0.05, *** P < 0.001.

## References and Notes

1. (available at https://idf.org/aboutdiabetes/what-is-diabetes/facts-figures.html).

2. E. A. Werner, J. Bell, J. Chem. Soc. Trans. 121, 1790–1794 (1922).

3. N. F. Wiernsperger, C. J. Bailey, Drugs. 58 Suppl 1, 31–39 (1999).

4. W. W. Wheaton et al., Elife. 2014 (2014), doi:10.7554/ELIFE.02242.

5. M. Y. El-Mir et al., J. Biol. Chem. 275, 223–228 (2000).

6. M. Foretz et al., J. Clin. Invest. 120, 2355–2369 (2010).

7. J. C. Geisler et al., Endocrinology. 154, 675–684 (2013).

8. K. Tatsushima et al., Biochim. Biophys. Acta - Mol. Basis Dis. 1867, 166013 (2021).

9. L. He, F. E. Wondisford, Cell Metab. 21, 159–162 (2015).

10. (available at https://www.proteinatlas.org/ENSG00000101194-SLC17A9/tissue).

11. S. Sakamoto et al., Sci. Reports 2014 41. 4, 1–10 (2014).

12. N. Hasuzawa et al., Yakugaku Zasshi. 141, 517–526 (2021).

13. C. Papandreou et al., Sci. Reports 2019 91. 9, 1–11 (2019).

14. K. C. Lin et al., Horm. Metab. Res. 44, 41–46 (2012).

15. S. Jain et al., Proc. Natl. Acad. Sci. U. S. A. 117, 30763–30774 (2020).

16. S. Jain et al., J.C.I. Insight. 6 (2021), doi:10.1172/J.C.I.INSIGHT.146577.

17. B. Thorens, Diabetologia. 58, 221–232 (2015).

18. X. Yang et al., Diabetologia. 62, 2106–2117 (2019).

19. N. Hashimoto et al., J. Biol. Chem. 262, 15026–15032 (1987).

20. Y. Zhang et al., Am. J. Physiol. Endocrinol. Metab. 302, 325–333 (2012).

21. M. Koike et al., Biochem. J. 283 (Pt 1), 265–272 (1992).

22. S. Keppens, H. De Wulf, Biochem. J. 240, 367–371 (1986).

23. S. Keppens et al., Br. J. Pharmacol. 105, 475–479 (1992).

24. S. Keppens, H. De Wulf, Biochem. J. 231, 797–799 (1985).

25. K. Enjyoji et al., Diabetes. 57, 2311–2320 (2008).

26. K. Marques Capiotti et al., Purinergic Signal. 12, 211–220 (2016).

27. D. L. Sparks et al., Recept. Clin. Investig. 1, e344 (2014), DOI:10.14800/RCI.344.

28. D. B. Buxton et al., Biochem. J. 237, 773–780 (1986).

29. X. Yang et al., Front. Endocrinol. (Lausanne). 10, 497 (2019). DOI: 10.3389/fendo.2019.00497

30. C. Wilcock, C. J. Bailey, Xenobiotica. 24, 49–57 (2009). http://dx.doi.org/10.3109/00498259409043220.

31. Y. Zhang et al., Front. Endocrinol. (Lausanne). 11, 341 (2020). DOI: 10.3389/fendo.2020.00341.

32. L. Ding et al., J. Biol. Chem. 286, 27027–27038 (2011).

33. W. Ma et al., J Biol Chem. 288, 15481–15494 (2013).

34. W. Ma et al., Biochem Pharmacol. 89, 99–108 (2014).

35. Y. Liu et al., J Biol Chem. 291, 1553–1563 (2016).

